# Immune landscape of a genetically-engineered murine model of glioma relative to human glioma by single-cell sequencing

**DOI:** 10.1101/2020.05.11.088708

**Authors:** Daniel B. Zamler, Takashi Shingu, Laura M. Kahn, Cynthia Kassab, Martina Ott, Katarzyna Tomczak, Jintan Liu, Yating Li, Ivy Lai, Cassian Yee, Kunal Rai, Stephanie S. Watowich, Amy B. Heimberger, Giulio F. Draetta, Jian Hu

**Affiliations:** The University of Texas MD Anderson Cancer Center Department of Genomic Medicine; The University of Texas MD Anderson Cancer Center Department of Cancer Biology; The University of Texas MD Anderson Cancer Center UT Health Graduate School of Biomedical Sciences; The University of Texas MD Anderson Cancer Center Department of Immunology; The University of Texas MD Anderson Cancer Center Department of Neurosurgery; The University of Texas MD Anderson Cancer Center Department of Melanoma Medical Oncology

**Keywords:** GBM, Quaking, QPP Model, Single Cell Sequencing, Immunotherapy, QKI

## Abstract

**Background:** This study focuses on the evaluation of intratumoral immune infiltrates of the *Qk/trp53/Pten* (QPP) triple-knockout mouse model of glioblastoma with the purpose of establishing its relevance compared to the human disease. This included an analysis at the single cell level, of immune cells composition in both spontaneous and implanted mouse tumors and of samples obtained from human tumor resections.

**Methods:** We analyzed a cohort of fifteen spontaneous QPP mice, nine implanted QPP mice and 10 glioma patients by standard immune profiling methods such as IHC. Of those, we analyzed three spontaneous QPP mice, three implanted QPP mice, and 10 glioma patients by single cell RNA sequencing.

**Results:** In the QPP samples we identified a predominantly myeloid cell population of monocytes, macrophages, and microglia, with minor populations of T, B, and NK cells. When comparing spontaneous and implanted mouse samples, we found that there were more neutrophil, T and NKT cells in the implanted model. In human samples, most of the lymphoid and myeloid compartments were immunosuppressive. While the complexity of the myeloid cell populations is preserved between the QPP model and the human disease, we observed significantly more immune-stimulatory populations in the implanted mouse model.

**Conclusions:** We found that, despite differences at the subpopulation level, the immunosuppressive nature of the immune system in glioma, is recapitulated in our mouse model. Although we observed differences in the proportions of immune infiltrates derived from the spontaneous and implanted mouse models and in LGG compared to GBM, the major constituents are present in all cases and these differences may be caused by various etiological or pathogenic influences on tumor-immune interactions.

**Key Points:**

- New GBM mouse model for immunotherapy
- Single cell profiling of the immune systems in human and mouse GBM

## INTRODUCTION

Glioblastoma (GBM) is the most frequent and deadliest primary brain tumor. For more than 40 years, chemotherapy, surgery, and radiotherapy have remained the standard-of-care treatment, and these approaches yield a dismal median survival of only 15 months. (Gao *et al*., 2013) With the advent of immunotherapy, a number of cancers previously recalcitrant to all therapies have been successfully treated and even cured. Unfortunately, immunotherapy in its current form has not been shown to exert a clinical benefit for most GBM patients, probably due to T-cell immune exhaustion, sequestration in the bone marrow, and an overwhelming immune-suppressive myeloid cell population. (Chongsathidkiet *et al*., 2018; Woroniecka *et al*., 2018)(De Groot *et al*., 2018) In GBM tumor-associated myeloid cells (TAMs), which likely originate from both tissue-resident microglia and monocytes recruited from the peripheral circulation, are immunologically heterogeneous, with the function of the various subpopulations unclear. This issue is further confused by lack of harmonization of nomenclature to describe TAM subpopulations. Notably, the TAM infiltrate of GBM remains understudied relative to infiltrating T cells, probably due to the success obtained by targeting T cells in other solid tumor types.

Currently, there is a paucity of spontaneous GBM models that enable preclinical studies of immunotherapeutic approaches. The GL261 mouse glioma system is one of the most commonly used models, although these tumors can grow in animals with an intact immune response this model lacks clonotypic diversity, is highly antigenic with a high tumor mutational burden, (Johanns *et al*., 2016) and is responsive to therapies that have failed in human clinical trials. (Genoud *et al*., 2018) To address the need for a preclinical model of GBM that can more faithfully recapitulate the human tumor environment, we characterized a murine model in which gliomas arise spontaneously by deletion of three common tumor suppressor genes in human gliomas: Quaking (*Qk* in mouse and *QKI* in human)(Verhaak *et al*., 2010), *trp53*(Cerami, Gao, Dogrusoz, Benjamin E Gross, *et al*., 2012) and *Pten*(Cerami, Gao, Dogrusoz, Benjamin E Gross, *et al*., 2012), the QPP model. *QKI* is mutated or deleted in around 34% of human glioblastomas (30% hemizygous deletions, 2% homozygous deletions, and 2% mutations). (Brennan *et al*., 2013) In addition, *QKI* downregulation through methylation of the *QKI* promoter was reported in 50 (20%) of 250 glioblastoma samples. (Chen *et al*., 2012) The significance of *QKI* in brain tumors is further highlighted by the observation that 90% of pediatric low-grade gliomas (angiocentric gliomas) contain *MYB-QKI* fusions that transform cells by concomitantly activating *MYB* and suppressing *QKI*. (Bandopadhayay *et al*., 2016; Qaddoumi *et al*., 2016) The QPP model uses an inducible cre-lox recombination system to delete the aforementioned genes under a *nestin* promoter that is expressed in neural stem cells, which generates tumors that are histopathologically similar to human GBM. Moreover, the spontaneously arising tumors are heterogeneous, and we have isolated and described murine tumors with features of the four described subtypes of human GBM demonstrating the concordance of this model and the human disease at a tumor cell level. (Shingu *et al*., 2017)

In this study, we aim to characterize the immune microenvironment of our previously established genetically-engineered mouse glioma model. Given the previously documented limitations of preclinical glioma models, heterogeneous immune infiltrates in glioma patients, and failure of immune therapies to provide broad and effective treatment, we aimed to immunologically characterize our murine glioma model for concordance with human gliomas.

We characterized the immune cell infiltration of a cohort of spontaneous and implanted QPP murine gliomas by immunohistochemistry (IHC), and single-cell RNA sequencing (scSeq) which offered a unique opportunity to dissect the immune infiltrates of murine and human gliomas while simultaneously determining the level of phenotypic concordance between the two. We found that tumors spontaneously arising in our murine glioma model recapitulated the heterogeneity and inter-patient variability of human gliomas demonstrated in this manuscript, carried a similar representation of immune cell repertoires compared to human glioblastoma, and therefore provide a suitable model for testing and triaging candidate immune therapeutics for patients with central nervous system (CNS) gliomas.

## IMPORTANCE OF THIS STUDY

This study lays the groundwork for future brain tumor research in two important ways. First, this is a resource for the brain tumor community for investigating immune infiltrates of glioma patients at a single cell level – this first of its kind. Second, we provide evidence that a newly established spontaneous and inducible mouse GBM model (QPP Model) recapitulates the constituents of the human immune response (predominantly myeloid with T, B, and NK cell components) as well as the complexity of the predominant immune population, myeloid cells, which include nine subtypes – monocytes, microglia, complement expressing microglia, macrophages, M0-like macrophages, M1-like macrophages, M2-like macrophages, dendritic cells, and neutrophils. This model provides a needed resource for immunocompetent studies of GBM in mice. While this model is beginning to gain interest and be used more broadly (Chen *et al*., 2019, 2020) this is the first comprehensive characterization of the immune compartment of the QPP model.

## Results

### Spontaneous and implanted QPP mouse GBM tumors mimic human GBM

We first confirmed the results of Shingu et al. demonstrating the concordance of the QPP model and human GBM from the perspective of tumor cells. (Shingu *et al*., 2017) To accomplish this, we generated glioma cell lines from spontaneously arising QPP tumors (**Supplemental Fig. 1a**) and these were able to form secondary tumors when implanted orthotopically into the striatum of C57BL/6J mice (**Supplemental Fig. 1e**). Both spontaneous and implanted QPP tumors showed necrotic areas (**Supplemental Fig. 1b, f**), invasive leading edge (**Supplemental Fig. 1c, g**), and a high proliferative index by Ki67 staining (**Supplemental Fig. 1d, h**), all key histopathological features of human GBM. Consistent with our previous observation, animals with spontaneous QPP tumors had a median survival of approximately 90 days (Shingu *et al*., 2017) (**Supplemental Fig. 2a**), whereas C57BL/6J mice orthotopically injected with QPP7, a cell line established from a spontaneous QPP tumor, had a median survival of approximately 45 days (**Supplemental Fig. 2b**).

We next performed immunohistochemistry analysis of the major immune constituents in 5 spontaneous and 5 implanted QPP tumors. We observed myeloid and lymphoid infiltrates, including T and Natural Killer (NK) cells, in both spontaneous and implanted tumors. (antibodies listed in **Supplementary Table** 1) The staining with antibodies directed at myeloid cell markers, such as Cd11b and arginase 1, was much more pervasive and intense relative to that of the lymphoid compartment in both tumor models, consistent with reported studies from human GBM the quantification of these stains and representative images are presented in **Figure 1** and **Supplementary Figure 2a,b**. There was no immune-cell infiltration detected in non-tumor-bearing control brain tissue isolated from C57BL/6J mice (**data not shown**), demonstrating that these immune populations are restricted to the pathogenic brain. These results were quantified and found to be statistically significant in that more myeloid and T-cells infiltrate in the implanted model than the spontaneous model and that there were significantly more myeloid cells than T-cells in both models (Hussain *et al*., 2006; De Groot *et al*., 2018) Together, our data indicate that both spontaneous and implanted QPP tumors exhibit histological characteristics and associated immune infiltrates similar to human GBM.

**Figure 1:**
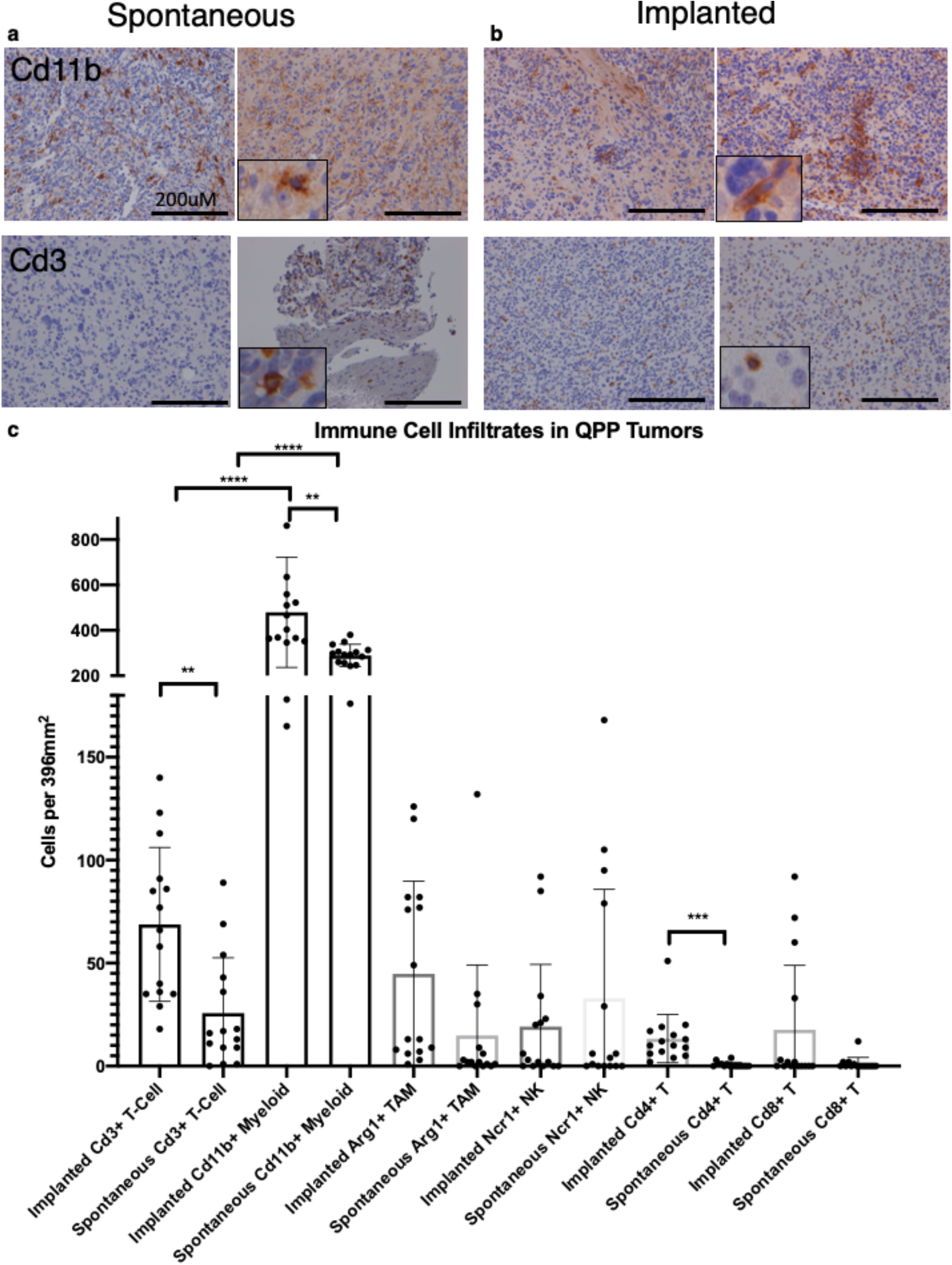
Immunohistochemistry (IHC) of immune markers confirming CIBERSORT analysis. **a**. Staining showing representative images of the brain of mice with no spontaneous or implanted tumor. **b**. Staining showing representative images of spontaneously arising QPP gliomas in mice when moribund. **c**. Staining showing representative images of implanted QPP gliomas in mice when moribund. Immune markers are displayed on the left: Arginase-1 (Arg1) for tumor-associated myeloid cells (TAM); lysosomal-associated membrane protein 1 (Lamp1) for myeloid cells; Cd3 for pan T cells; Cd4 for T-helper cells; Cd8 for cytotoxic T-cell lymphocytes (CTL); Natural-killer-cell receptor 1 (Ncr1) for natural killer cells. Large images shown at 20 x magnification. Insert images shown at 4 x; scale bars = 200 µM; M indicates mouse number; DPB means Days Post Birth; DPI means Days Post Injection.

### Single-cell sequencing reveals that the components and complexity of human disease are represented in the QPP model

Approaches like IHC use single markers or limited marker profiles several to define cell populations, but these techniques may be unable to account for all differences that may exist in the immune cell populations from diverse species. (Mestas and Hughes, 2004, Shay *et al*., 2013, Yue *et al*., 2014) Thus, to investigate the immune compartment of GBM at the single-cell level, we conducted experiments using the 10x (10x Genomics; 6230 Stoneridge Mall Road, Pleasanton, CA 94588) single-cell sequencing platform utilizing mouse and human GBM tissues. Specifically, we evaluated either freshly isolated or previously frozen samples, and then looked at a batch corrected cohort of CD45^+^-enriched populations of immune cells from spontaneous (n=3) and implanted (n=3) QPP tumors, as well as from a cohort of human tumors (n=10). (basic mutational and treatment summary in **Supplementary Table 2**) Furthermore, we compared batch corrected CD45^+^ cells isolated from fresh tumor-free mouse brains with those from frozen tumor-free mouse brains and found them to be comparable regarding their myeloid and lymphoid constituents (**Supplemental Fig. 3**). For spontaneous tumors, by clustering using 9 principal components as chosen by elbow plot, to reveal the major constituents of the immune infiltrates present in the mouse model [resolution = 0.1] we identified four subtypes (**Fig. 2a**): macrophages, microglia (**Supplemental Fig. 6b**), neutrophils; (**Supplemental Fig. 6c**), myeloid antigen-presenting-like cells (APCs; **Supplemental Fig. 6d**). T, B and NK cells were also detected in these samples, but their abundance was not sufficiently high to meet the threshold required for the analysis to assign independent clusters (**Supplemental Fig. 6e, f, g**). All clusters were defined based on the three representative markers shown in the supplementary figures, as well as a variety of others, that were selected from the top differentially expressed genes for that cluster. The full list of genes upregulated for each cluster can be found in **Supplementary Table 3**. The same analysis performed on the cohort of implanted QPP tumors identified 5 clusters (**Fig. 2b**): microglia/macrophages (**Supplemental Fig. 8b**), neutrophils (**Supplemental Fig. 8c**), APCs (**Supplemental Fig. 8d**), lytic myeloid cells (**Supplemental Fig. 8e**), and T, B and NK cells (**Supplemental Fig. 8f, g, h**). The full list of genes upregulated for each cluster can be found in **Supplementary Table 4**. These cell populations are consistent with our IHC and are also congruent with reported human data (Chen and Hambardzumyan, 2018). We compared the implanted and spontaneous models by combining them into one dataset (**Figure 2c**) and found five clusters, neutrophils (**Supplemental Fig. 10b**), microglia/macrophages (**Supplemental Fig. 10c**), myeloid APC-like cells (**Supplemental Fig. 10d**), lytic myeloid cells (**Supplemental Fig. 10e**) and T and NK cells, importantly a population of B cells was also present although not a cluster (**Supplemental Fig. 10f, g, h**). The full list of genes upregulated for each cluster can be found in **Supplementary Table 5**. Notably, our comparison uncovered a higher proportion of neutrophil and T/NK cell infiltrates in implanted than in spontaneous QPP tumors, which may indicate the different trajectories of tumor progression between the two models (i.e., the spontaneous tumors presumably start from one or a few cells rather than the 50,000-cell starting point with the implanted tumors). We also observed slight variations among different mice (some mice have more neutrophils than others), which recapitulates the heterogeneous nature of the human disease (**Supplemental Fig. 13a, b, c**). Importantly, this heterogeneity was not caused by technical variation, which was found to be minimal by examining two technical replicates (**data not shown**). In the cohort of human glioma samples, we identified seven clusters (**Figure 2d**): T cells (**Supplemental Fig. 12b**), neutrophils (**Supplemental Fig. 12c**), NK cells (**Supplemental Fig. 12d**), microglia or macrophages that are APC-like (**Supplemental Fig. 12e**) microglia or macrophages that are polarized (**Supplemental Fig. 12f**) and B cells (**Supplemental Fig. 12h**), which was consistent with the clusters identified in the spontaneous and implanted QPP models. Microglia and macrophages are present across all samples (**Supplemental Fig. 12g**). The full list of genes upregulated for each cluster can be found in **Supplementary Table 6**. The available mutational profile for all patients enrolled is available in **Supplementary Table 7**

**Figure 2:**
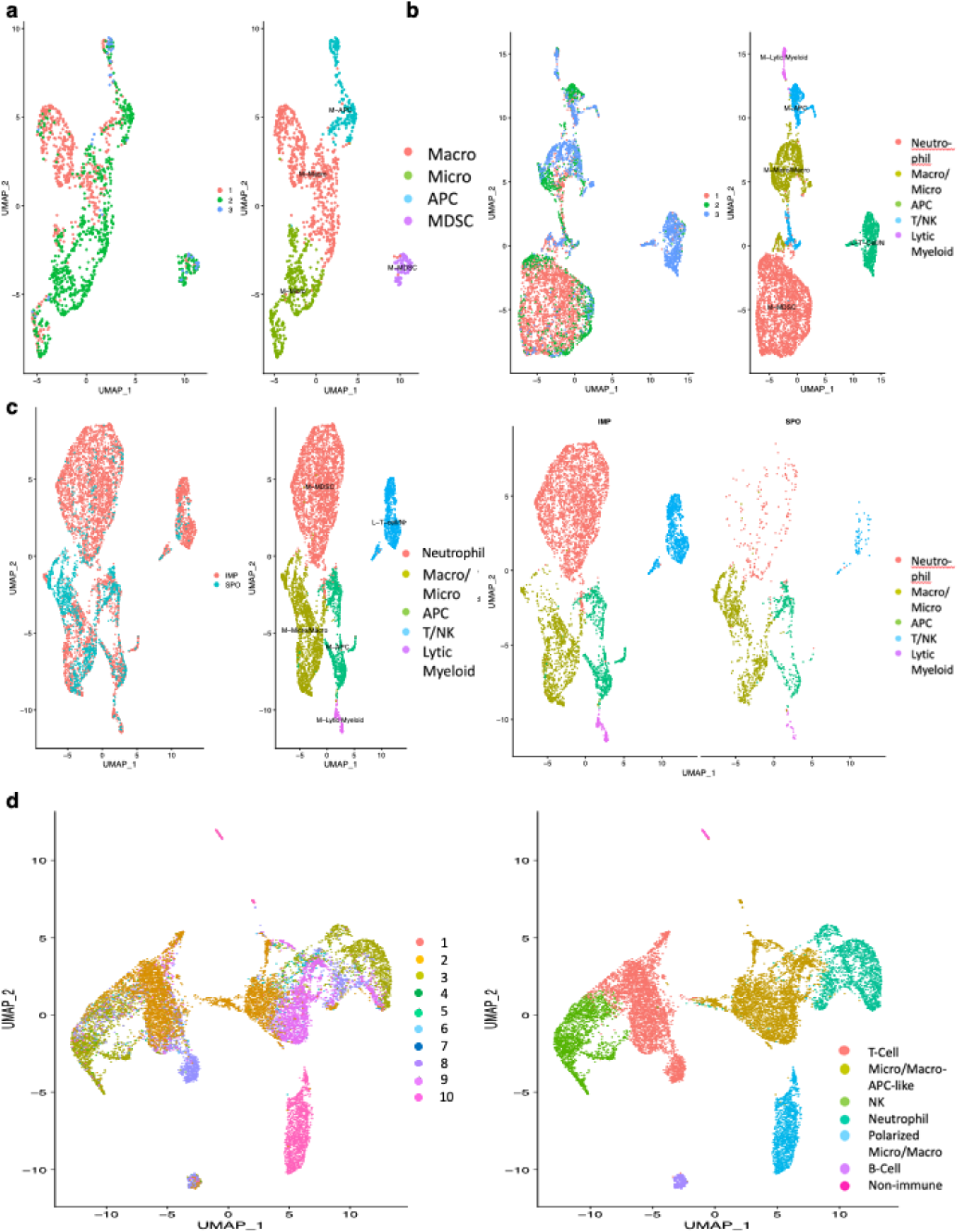
Clustering to reveal major immune constituents. Resolution (0.1) **a**. Clusters revealed for n=3 spontaneous QPP mice are macrophages, microglia, antigen-presenting-like cells (APCs), and neutrophils with 9 PC’s determined by elbowplot **b**. Clusters revealed for n=3 implanted QPP7 mice are neutrophils, microglia/macrophages, T cells/NKs, APCs, lytic myeloid with 9 PC’s determined by elbowplot **c**. Comparison of the implanted and spontaneous QPP tumor clusters shows neutrophils, microglia/macrophages, T cells/NKs, APCs, lytic myeloid with 9 PC’s determined by elbowplot **d**. Clusters revealed for n=10 glioma patients are T cells, APCs, NKs, neutrophils, polarized microglia/macrophage, B cells and non-immune. with 15 PC’s determined by elbowplot

Importantly, our human population was made up of 10 patients that were identified as GBM by MRI. Upon collection of the patient information, after pathologist consult, we learned that three of our patients were actually lower grade glioma (LGG) with one patient being diagnosed with oligodendroglioma (grade II) and two of our patients being diagnosed with anaplastic astrocytoma (grade II). We compared the major immune constituents [resolution = 0.1] between these two groups and found one myeloid population and one T-cell population that were over-represented in the LGG patients (**Supplemental Fig. 14a,b**). Three defining markers (differentially expressed genes) for the myeloid population were C1QA, a component of complement involved in immune response, MRC1, a scavenger receptor that is a marker for “M2-like” macrophages, and CCL3, a chemokine involves in macrophage inflammation response (cluster 3 in **Supplemental Fig. 14c)**, while two defining markers (differentially expressed genes) for the T cell population were CXCR4, a chemokine receptor associated with CD4 function in T cells, IL7R, the receptor for interleukin 7 that is important for VDJ recombination of T-cells (cluster 4 in **Supplemental Fig. 14d)**. This indicates that cells of these immune subtypes are under-represented in GBM compared to LGG, however none of these differences impacted our downstream analyses. The differences, however, do warrant further investigation. For consistency, the same markers (or homologs) were used to designate these populations regardless of the species, except that CD24a and ITGAX were used interchangeably to label neutrophil clusters, and Nktr and HCST were used to label NK clusters in mice and humans, respectively.

We next performed subtype clustering to determine the minor immune constituents [resolution = 0.65] and to identify subpopulations among the major immune-cell clusters (**Fig. 3a, b**). In our mouse dataset we found populations of Cxcl12^hi^Cd14^hi^ neutrophils, Cd14^lo^ neutrophils, Il-1b^hi^ neutrophils, Cd11c^+^Csf1r^+^Cx3cr1^+^ macrophages, Il-1b^lo^ neutrophils, Treg/NKT, Tgfb1^+^ macrophages, MHCII^+^ APC’s, Ifn gamma responsive monocyte/macrophage, metabolically active monocyte/macrophage, Cd8^+^ T cells, microglia, metabolically active lymphoid, B cells, dendritic cells, and pDCs. In our human dataset we found populations of IL7R^+^ T or lymphoid, APOE^+^APOC1^+^ macrophages, CD8^+^ T cells, MRC1+ macrophages, SPP1+ macrophages, LYZ^hi^ neutrophils, NKG7^hi^ ILCs, PRF1^hi^ ILCs, DCs, metabolically active lymphoid, FN1^+^ macrophages, B cells, IFNg^+^ CD8^+^ T cells, LYZ^lo^ neutrophils, HSP/ribosome protein enriched lymphoid cluster, metabolically active macrophages, minor myeloid population, and two sets of non-immune populations which suggests we successfully captured cells with a range of CD45 expression. The full list of genes upregulated for each cluster can be found in **Supplementary Table 8 (mice) or Supplementary Table 9 (human)**. We identified consistency within the lymphoid compartment with regard to T cells marked by Cd3d, e, g, in the mouse (CD3D, E, G; in the human); NK cells - Klrd1, Nkg7, Nktr (KLRD1, NKG7, NKTR in human); and B-cells - Cd79a, Cd79b, Ms4a7 (CD79A, CD79B, MS4A7 in human) (**Figure 4a, b**). By separating the myeloid compartment into subsets, we could identify representative markers such as Cd14 in mouse (CD14 in human) – monocytes; Grn (GRN in human) – microglia; - C1qa (C1QA in human) – complement-expressing microglia; Itgam (ITGAM in human) – macrophages; Pirb (LILRB1 in human) – M0-like macrophages; Il1b (IL1B in human) – M1-like macrophages; Mrc1 (MRC1 in human) – M2-like macrophages; Itgax (ITGAX in human) – dendritic cells; and S100a8 (S100A8 in human) – myeloid-derived suppressor cells. We found that each of these myeloid subtypes were detected in both the mouse and human datasets, and given the spectrum nature of myeloid cell polarization it is unsurprising that many of these markers show up across multiple myeloid subtypes (**Fig. 5a, b**). Taken together, single-cell analysis of the immune infiltrate of murine and human GBM tissues was consistent with our histopathological analysis, supporting the idea that both the spontaneous and implanted QPP murine model tumors faithfully recapitulate the immune compartment of human GBM. Given the difficulty in mapping immune subset populations between mouse and humans we aimed to abstract this information into relevant gene ontological pathways that were conserved between mouse and human immune cells and what populations upregulate these ontologies.

**Figure 3:**
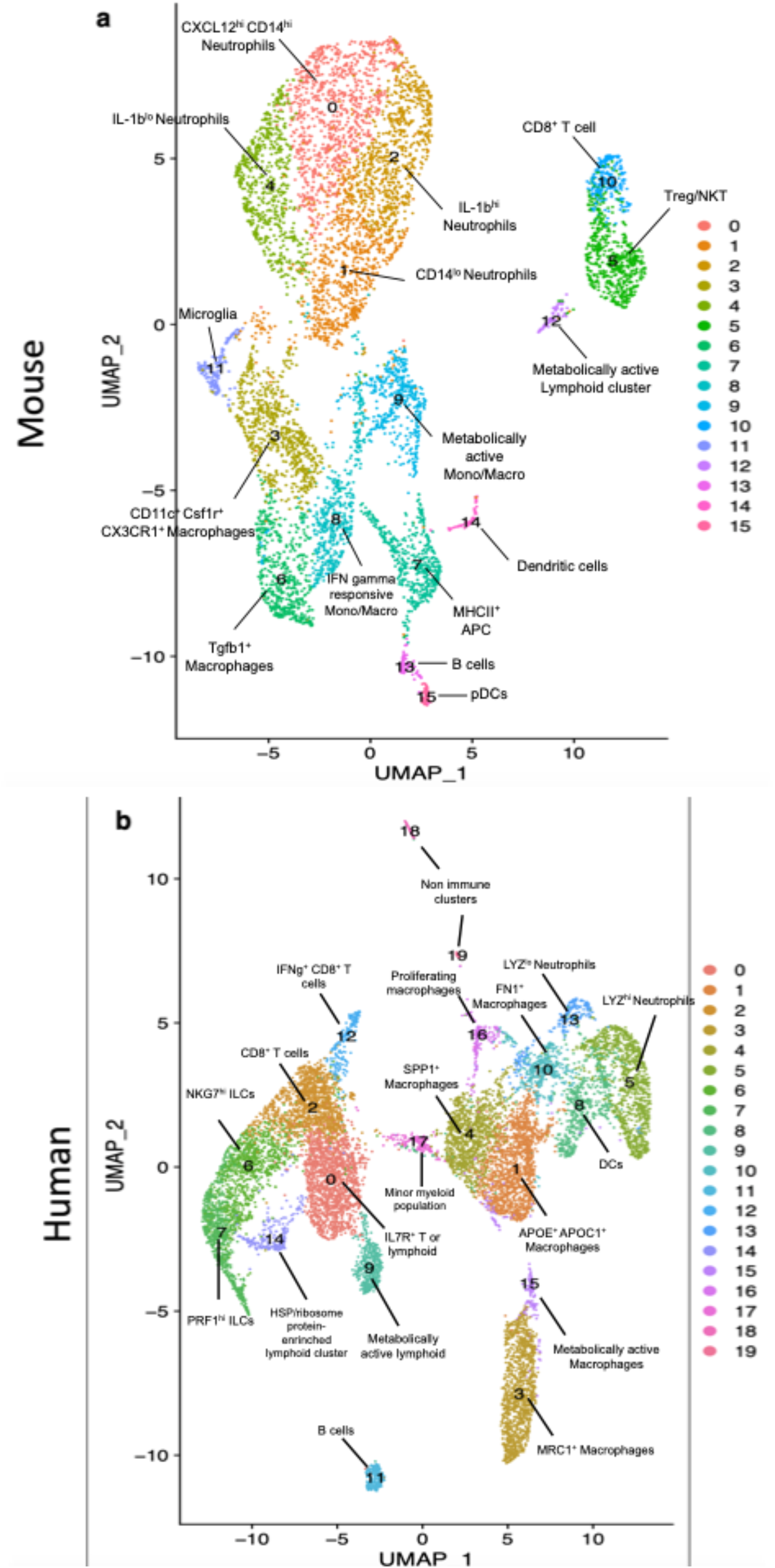
Clustering to reveal subtypes. Resolution (0.65) **a**. Combined mouse spontaneous and implanted QPP tumor dataset with 9 PC’s determined by elbowplot. **b**. Aggregated human GBM dataset with 15 PC’s determined by elbowplot.

**Figure 4:**
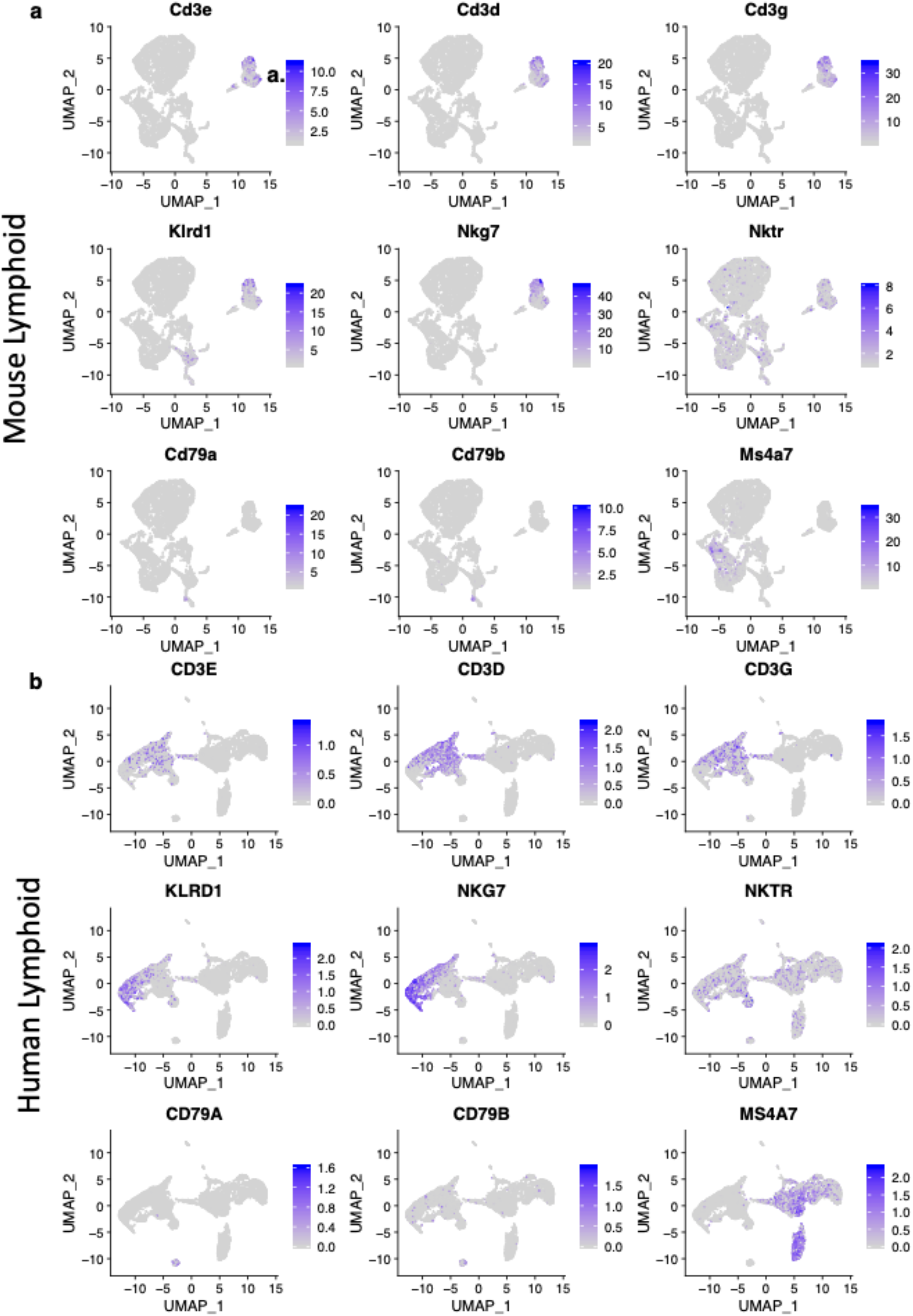
Lymphoid cell subtypes of human GBM are contained in the QPP model. Resolution (0.65) clustering to reveal lymphoid subtypes in the QPP mouse model tumor or human GBM. **a, b**. Representative markers for the following populations: Cd3d, e, g (CD3D, E, G) – T cells; Klrd1, Nkg7, Nktr (KLRD1, NKG7, NKTR) – NK cells; Cd79a, Cd79b, Ms4a7 (CD79A, CD79B, MS4A7) – B cells.

**Figure 5:**
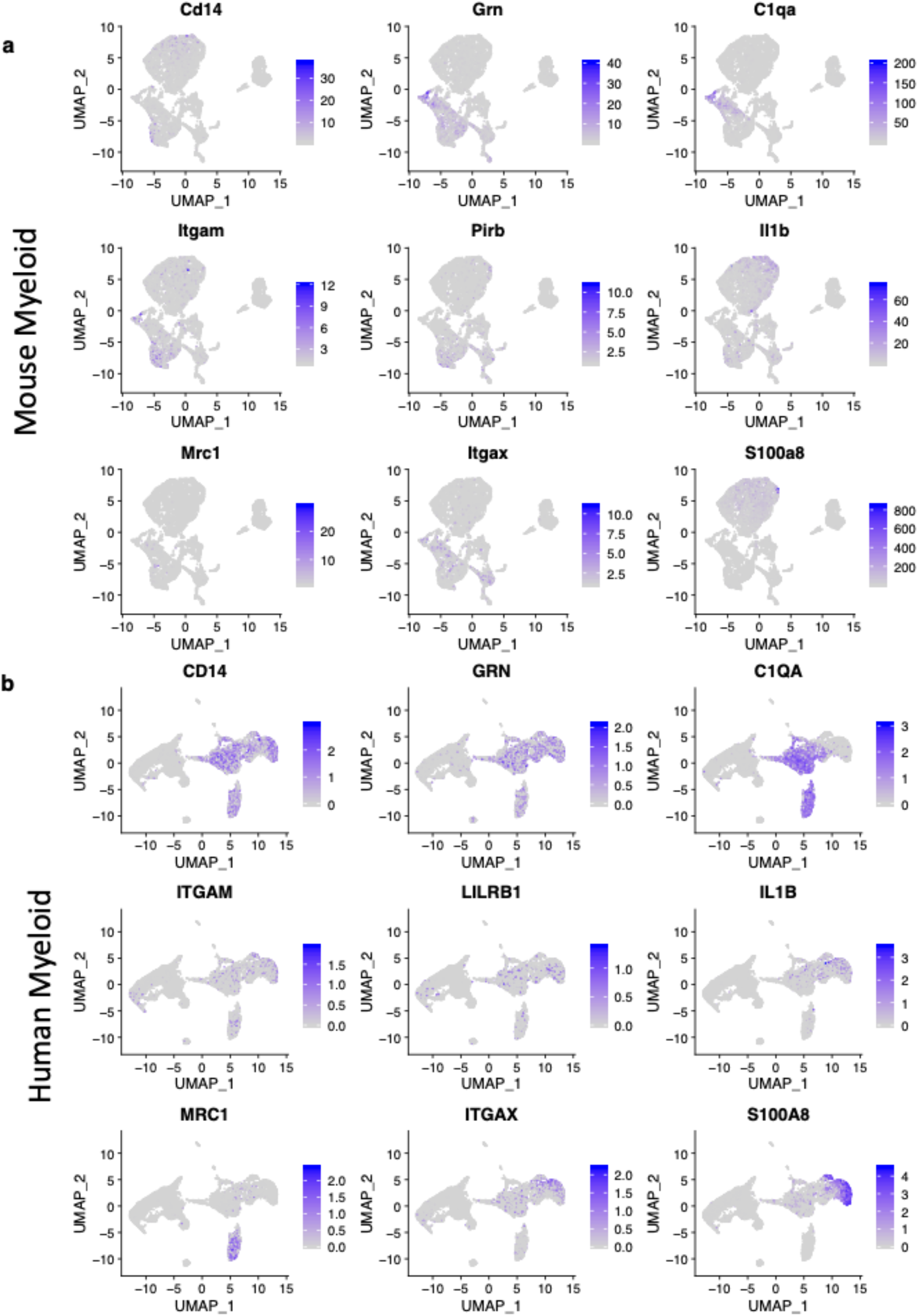
Myeloid cell subtypes of human GBM are contained in the QPP model. **b**. Resolution (0.65) clustering to reveal myeloid subtypes in the QPP mouse model tumor or human GBM. **a, d**. Representative markers for the following populations Cd14 (CD14) – monocyte, Grn (GRN) – microglia, - C1qa (C1QA) – complement-expressing microglia, Itgam (ITGAM) – macrophages, Pirb (LILRB1) – M0-like macrophages, Il1b (IL1B) – M1-like macrophages, Mrc1 (MRC1) – M2-like macrophages, Itgax (ITGAX) – dendritic cells, S100a8 (S100A8) – myeloid-derived suppressor cells. Given the spectral nature of myeloid differentiation and polarization many of these markers will be present in multiple subsets.

In order to quantify the diversity of immune species in all of our samples we calculated the Shannon Diversity Index (SDI) for each of our datasets. The results of these analyses are listed in **Supplementary Table 10** and, briefly, show that the human dataset is the most diverse (9.3 SDI) and the spontaneous dataset is the least diverse (7.16 SDI). The implanted dataset falls in the middle (8.54 SDI) and the combined mouse dataset is similar to the implanted alone (8.76 SDI). Due to the lack of sensitivity associated with the SDI, we furthermore performed a Chi-squared test of heterogeneity on our samples and also found them to be statistically significantly different

### Gene ontology comparison shows conserved immune pathways

Given the large list of differentially expressed genes generated by each cluster at higher resolution and the difficulty in performing gene mapping from one species to another, we sought to identify high-level pathways that are conserved between our murine models and the human disease. To this end we performed Gene Ontology analyses as previously carried out to determine what high-level pathways might be conserved in mouse and human. (Chen, 2020) All gene ontologies can be found in (**Table 1**) if none are listed, none passed the threshold of a 0.05 > p-value.

**Table 1:** Conserved GO Pathways from Immune Isolates of QPP Knockout Tumors and Human GBM. We took the full list of positively differentially expressed genes for each cluster at the subtype resolution [0.65] we performed GO pathway analysis on this full list for each cluster and catalogued the top 10 gene ontologies for each cluster in biological processes, molecular function, and cellular component. Numbers in green denote the human clusters upregulating the GO pathway while those in purple represent the mouse cluster that has the GO pathway upregulated.

## Discussion

The limited availability of animal models that faithfully recapitulated the pathogenesis, histopathological characteristics and disease development of glioblastoma is a likely significant contributor to the poor clinical success rate of candidate drugs tested in this disease setting. Here we characterize the QPP model in both the spontaneous and implanted form and their immune infiltrates with relative concordance to the human disease. We encourage others to characterize other available models, as there is a paucity of models in general for gliomas, and furthermore for studies that require an intact immune system. (Implantation of GL261 cells into C57BL/6J mice is used almost exclusively) The GL261 cell line was generated by injecting methylcholanthrene into the brain of C57BL/6J mice, and divergent clones subsequently derived from it have been used by various groups. (Szatmári *et al*., 2006) GL261 tumors harbor activating *KRAS* mutations and express high MHC-1 protein levels but have relatively limited expression of other immune antigens, including MHC II, B7-1, and B7-2, resulting in a limited, immunogenic phenotype. As previously described, our inducible QPP mice develop heterogeneous tumors with features similar to all four human GBM subtypes recapitulated in different mice, and it is thus reasonable to speculate that further characterization of our model, as well as of human GBM tumors, may reveal that the QPP model offers an opportunity to model immunogenically diverse populations similar to the human disease as shown by our single cell sequencing dataset. Other spontaneous models of GBM include the sleeping beauty transposase-mediated (Ohlfest *et al*., 2005) and virally-induced systems (Miyai *et al*., 2017) that rely on strong mutagenic drivers not common in human GBM, such as mutant *KRAS*. In contrast, our spontaneously-arising GBM model is induced by deletion of three key tumor-suppressor genes that are frequently mutated/deleted in human GBM patients. (Verhaak *et al*., 2010; Gao *et al*., 2013)

We found that along with histological features of human GBM, our model closely approximates the immune infiltrates of the human GBM microenvironment by histopathological analysis, and scSeq. Although we observed slight variations among data sets (e.g., T cells lower in mice than humans by scSeq), a predominantly myeloid cell immune compartment was confirmed by both analyses in both species. Furthermore, subtype analysis of the myeloid compartment demonstrated similar complexity and populations between mouse and human glioma cohorts, although they appear to be driven by different transcriptomic programs. We observed an enrichment of neutrophils, T cells, and NK cells in implanted relative to spontaneous QPP tumors, which might be influenced by the microenvironmental response to the mechanical implantation procedure and the different tumor progression trajectories of these tumors. The spontaneous model presumably starts from one or a few cells which is transformed by cre-lox recombination and then must progress and become aggressive, whereas the implanted tumor starts with a bolus of 50,000 cells that have survived the selective pressure of serial passaging in vitro. Overall, our data indicate that the spontaneously arising GBM model may be a closer approximation of human GBM and more suitable for testing certain types of immunotherapy requiring an intact immune system.

Although our number of patients and mice is relatively small (n=6 mice; n=10 patients) compared to other GBM datasets, (Cerami, Gao, Dogrusoz, Benjamin E. Gross, *et al*., 2012) ours has the advantage of being selected specifically for immune cells while other datasets only contain them as an ancillary point to the tumor cells. We furthermore retain the major immune populations in our samples consistent with the literature previously reported (*i*.*e*. macrophages). (Chen and Hambardzumyan, 2018) It has been shown previously that treatment in sample preparation of the immune cells can differentially affect immune cell populations, and all samples we analyzed were subjected to a cleanup process after being reduced to a single-cell suspension. Furthermore, a portion were also frozen at -80*C for a time. These processes undoubtedly affect certain immune-cell populations differently and may induce a sort of “selection” in our downstream analyses. We controlled for this in the experiments presented here, as all mouse and human tissues were treated in a consistent fashion and major populations were conserved between previously frozen and freshly isolated samples (**data not shown**).

Our human dataset showed a higher proportion of T-cell and NK-cell infiltration than would be expected based on previous studies.(Chen and Hambardzumyan, 2018) This is potentially due to our procedure, which involved freezing, and seems to affect human and mouse tumor samples differently. Indeed, Kadic et al. have reported that the freezing process has a more negative effect on myeloid cell populations than lymphoid populations.(Kadić *et al*., 2017) Nevertheless, although our human dataset has higher lymphoid immune infiltrates than expected, we can see that most of these come from the sample from patient 2 whose profile, along with the others can be found in **Supplementary Table 7**, whereas the other 9 patients’ samples exhibited myeloid cells more consistently. When looking at the subset of immune cells in our mouse and human datasets, we found that both the constituents (T cells, B cells, NK cells, and myeloid cells) and the complexity of these compartments were effectively recapitulated in the QPP mouse model.

Further confirming our observations in both mouse and human, a recent back-to-back publication in Cell (Friebel *et al*., 2020; Klemm *et al*., 2020) that patients have a highly variable immune infiltrating microenvironment composed of myeloid, neutrophil, and T-cell components with smaller proportions of other immune populations present. While these observations are not the main point made in these papers they confirm and support our findings of both the human disease and mouse model. Friebel et al performed CyTOF analysis providing a protein level view of the immune populations and how their infiltrates vary based on metastasis and IDH status while Klemm et al investigated the transcriptomic changes that occur with different tumor origin and IDH status.

Due to the complexity of mapping the mouse genome to the human genome (e.g. differences in capitalization, differences in gene names, homologs, etc.), we sought to compare the immune subtypes of the human and mouse glioma datasets with gene ontologies [resolution = 0.65]. As expected, we observed a large proportion of gene ontologies revolving around canonical immune functions, such as immune response, defense response and regulation of immune system processes. We also found less expected upregulated gene ontologies peptide and amide synthesis in the T-cell compartment and peptide and amide binding in the myeloid compartment which suggests a signaling mechanism that warrants further investigation.

Given all of these differences, there is concordance between the QPP mouse model and the human disease. While there are small populations that exist in the mouse dataset that are of a more immunostimulatory nature than observed in the humans both of our datasets are dominated by immunosuppressive myeloid populations. Many of the same populations were observed in both systems although driven by different cellular programs.

Our work provides further insight at single cell level for therapeutic studies targeting the immune system in GBM. We believe our work will allow better preclinical modeling of the human disease in labs across the globe, which hopefully will promote better translation of novel therapeutics to the clinic.

## Acknowledgements

The authors acknowledge Angela K. Deem Ph.D. and David M. Wildrick, Ph.D., for their editorial support. In memory of Sherwin Elliot Zamler.

## Author Contributions

Experimental design and/or implementation: DBZ, CK, MO, KT, JL, YL, IL, CY, KR ABH, GFD, JH

Analysis and interpretation of the data: DBZ, JH, ABH, GFD

Writing of the manuscript: DBZ, ABH, GFD, JH

## STAR METHODS

### Lead contact

Further information and requests for resources and reagents should be directed to and will be fulfilled by the Lead Contact, Jian Hu 1818 East Rd. 3SCR5.3612, Houston, TX 77054; P: 713-794-5238; F: 713-792-9636;

### Materials Availability

- Cell Lines – QPP7 cell line was generated in 2017 at MDACC in the Hu laboratory, as the cells are primary, they have not been banked, tested, or authenticated, however the cells are available for sharing with the community upon request.
- Reagents – This study did not generate any unique reagents
- Mouse lines – This study did not generate and unique mouse lines

### Data and Code Availability

- Dataset – The single cell sequencing dataset has been deposited in the GEO database under the accession number GSE147275 with access token opejqqwmtvwxhwb
- Code – Code used to generate figures and analyze data is available at https://github.com/zamlerd/Single_Cell_Sequencing

## Experimental Model and Subject Details

### Mouse Models

The QPP spontaneous glioma model exists on a mixed background and is maintained in the Hu laboratory at MD Anderson Cancer Center in the Department of Cancer Biology, Frozen Sperm have also been deposited in the MD Anderson Mouse Transgenics Core

### Cell Lines

Cell Lines - Cell line was generated in 2017 at MDACC in the Hu laboratory, as the cells are primary, they have not been banked, tested, or authenticated, however the cells are available for sharing with the community upon request.

### Patient Data

Patient information with sex, genomic information as well as site of resection etc. are available in **Supplementary Table 7**.

## KEY RESOURCES TABLE

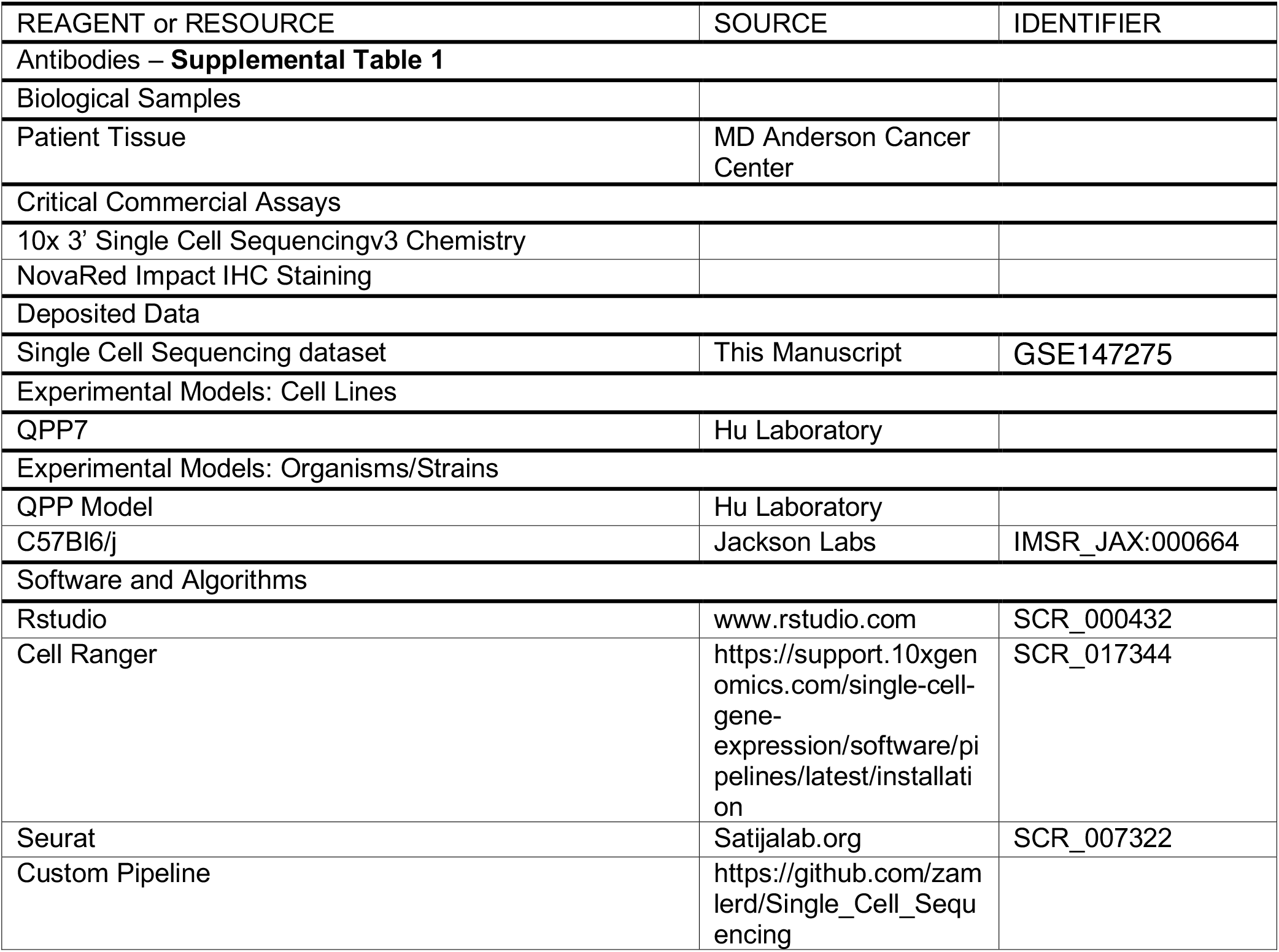

### Cell lines

The QPP7 cell line was cultured in the commercially available NeuroCult Basal Medium (Mouse & Rat; Cat # 05700; Stemcell Technologies Inc., Cambridge, MA) with the NeuroCult Proliferation Supplement (Mouse & Rat; Stemcell Technologies, Inc.) added. Cells were cultured as spheres and subcultured every 2-3 days at a 1:10 ratio with Accutase (Innovative Cell Technologies, Inc.; San Diego, CA) as the dissociation solution.

### Murine glioma models

All manipulations were performed with Institutional Animal Care and Use Committee approval at The University of Texas MD Anderson Cancer (MD Anderson). To trigger the spontaneous QPP gliomas, tamoxifen was dissolved in corn oil at the concentration of 10 mg/ml and injected subcutaneously in a total volume of 20 uL in P7 and P8 mice to induce glioma development in the Nes-CreERT2; Qk^L/L^; Pten^L/L^; Trp53^L/L^ background. The mice were monitored for neurological symptoms or other signs of ill health every other day and were euthanized and necropsied when moribund. To implant QPP gliomas, cells were cultured as described above until the time of surgical implantation. The QPP cells were dissociated with Accutase (Innovative Cell Technologies, Inc.) for 5-10 minutes at room temperature, the Accutase was neutralized by dilution with medium, and the cells were pelleted by centrifugation. Automated cell counting was performed and the cells were resuspended at a concentration of 2.5 × 10^4^ cells per uL. Mice were anesthetized with a combination of ketamine and xylazine and 5 × 10^4^ QPP cells were implanted at the stereotactic coordinates of +0.5 mm forward and +2 mm lateral right from the bregma at a depth of 3mm. After the anesthesia was reversed with atipamezole, the mice were monitored until signs of tumor burden appeared, at which point they were euthanized by transcardial perfusion with Tyrode’s solution (Sigma-Aldrich, Inc., St. Louis, MO.). Their brains were removed and fixed in paraformaldehyde.

### Immunohistochemistry

Tissues were embedded in paraffin, serially sectioned on a microtome at 5 uM, and stained with hematoxylin and eosin. Specifically, sections on microscope slides were stained with freshly filtered hematoxylin for 30 seconds and then with eosin for 15 seconds before dehydration in two 1-minute washes of 95% ethanol followed by three 1-minute washes in 100% ethanol, and finally, three quick rinses in xylene before application of cover slips to slides. For antibody staining, tissue sections were baked for 1 hour at 60 °C before being washed three times in xylene for 5 minutes, followed by washes in 100%, 95%, 70%, and 50% ethanol and then tap water. Antigen retrieval was performed using a Biogenix easy-retrieval microwave set at 95°C for 10 minutes in sodium citrate buffer at pH 6.0. Slides were then washed in PBS for 5 minutes before being blocked with 3% BSA at room temperature for 1 hour. Slides were then incubated overnight at 4°C at the dilutions listed **in Supplementary Table 1**. The next morning, slides were washed three times in PBS with 0.1% Tween for 5 minutes before incubation with the appropriate HRP-conjugated secondary antibody for 1 hour at room temperature. Slides were then washed three times in PBS for 5 minutes before being developed using the NovaRED chromagen incorporation kit. Cover slips were then applied to the slides in aqueous mounting medium. Antibody catalog numbers and dilutions and RRIDs are found in **Supplementary Table 1**.

### Single-cell sequencing

Mice with implanted QPP tumors were perfused with Tyrode’s solution before the tumors were reduced to a single-cell suspension and frozen in Bambanker cell-freezing medium (Wako Chemicals USA, Inc., Richmond, VA) at -80 °C. Gliomas were mechanically dissociated with scissors while suspended in Accutase solution (Innovative Cell Technologies, Inc.) at room temperature and then serially drawn through 25-, 10- and 5-mL pipettes before being drawn through an 181/2-gauge syringe. After 10 minutes of dissociation, cells were spun down at 420 x g for 5 minutes at 4 °C and then resuspended in 10 mL of a 0.9N sucrose solution and spun down again at 800 x g for 8 minutes at 4 °C with the brake off. Once sufficient samples were accumulated to be run in the 10x pipeline (10x Genomics; 6230 Stoneridge Mall Road, Pleasanton, CA 94588), cells were then thawed and resuspended in 1 mL of PBS containing 1% BSA, for manual counting. Cells were then stained with the CD45 antibody (BD Biosciences, San Jose, CA, cat #: 555482) at 1:5 for human or (Tonbo Biosciences, San Diego, CA, cat #: 50-0454-U100) at 1:10 for mice for 20 minutes on ice. Samples had Sytox blue added just before sorting so that only live CD45+ cells would be collected. Cells were then sorted in a solution of 50% FBS and 0.5% BSA in PBS, spun down, and resuspended at a concentration of 700-1200 cells/uL for microfluidics on the 10x platform (10x Genomics). The 10x protocol, which is publicly available, was followed to generate the cDNA libraries that were sequenced. (https://assets.ctfassets.net/an68im79xiti/2NaoOhmA0jot0ggwcyEKaC/fc58451fd97d9cbe012c0abbb097cc38/CG000204_ChromiumNextGEMSingleCell3_v3.1_Rev_C.pdf)

The libraries were sequenced on an Illumina next-seq 500, and up to 4 indexed samples were multiplexed into one output flow cell using the Illumina high-output sequencing kit (V2.5) in paired-end sequencing (R1, 26nt; R2, 98nt, and i7 index 8nt) as instructed in the 10x Genomics 3’ Single-cell RNA sequencing kit.

The data were then analyzed using the cellranger pipeline (10x Genomics) to generate gene count matrices. The mkfastq argument (10x Genomics) was used to separate individual samples with simple csv sample sheets to indicate the well that was used on the i7 index plate to label each sample. The count argument (10x Genomics) was then used with the expected number of cells for each mouse or patient. For the mice, the expected cell numbers were 10,000, whereas for patients, the numbers varied between 2,000 and 8,000. Mice were aligned with the mm10 genome, and humans were aligned with GRCh38. The aggr argument (10x Genomics) was then used to aggregate samples from each condition (spontaneous QPP, implanted QPP, and patient) for further analysis. Once gene-count matrices were generated, they were read into an adapted version of the Seurat pipeline(Butler *et al*., 2018; Stuart *et al*., 2019) for filtering, normalization, and plotting. Genes that were expressed in less than three cells were ignored, and cells that expressed less than 200 genes or more than 2500 genes were excluded, to remove potentially poor- and high-PCR artifact cells, respectively. Finally, to generate a percentage of mitochondrial DNA variability and to exclude any cells with more than 25% mitochondrial DNA (as these may be doublets or low-quality dying cells), cells were normalized using regression to remove the percent mitochondrial DNA variable via the scTransform (Hafemeister and Satija, 2019) command. Next, the cell clusters were identified and visualized using SNN and UMAP, respectively, before generating a list of differentially-expressed genes for each cluster. A list of differentially-expressed genes was generated to label our clusters at low resolution (0.1). These clusters’ labels were based on at least three differentially-expressed genes, and violin plots were generated to show the relative specificity to the cluster.

Identification of the clusters was as follows: Neutrophils: CD24a, S100a8, and S100a9; antigen-presenting cells: CD74 (MHCII), H2-Eb1, and H2-Aa; T cells: Cd3d, Cd3e, and Cd3g; and microglia and macrophages: Cd68, Cx3cr1, and Tmem119. For analyses performed on the combination of implanted and spontaneous QPP models, we joined the datasets (using the FindIntegrationAnchors command to determine genes that can be used to integrate two datasets—after the determination of the Anchors used the IntegrateData command) with the aforementioned anchors to combine our two datasets. Data were then normalized using the scTransform command, which uses regression analysis to remove the percentage of mitochondrial DNA from each cell. Datasets were then processed for principal component analysis (PCA) with the RunPCA command, and elbow plots were printed with the ElbowPlot command in order to determine the optimal number of PCs for clustering. Datasets then were submitted to cluster analysis with RunUMAP and FindNeighbors commands before FindClusters was run with either 0.1 or 0.65 resolution for low- and high-resolution clustering, respectively. Differentially-expressed genes were identified using cutoffs for min.pct = 0.25 and logfc.threshold = 0.25. Plots were generated with either the DimPlot, FeaturePlot or VlnPlot commands.

### Gene Ontology Analysis

Gene ontology analyses were performed as previously published and publicly available code adapted to our dataset. (Chen, 2020)

---

**Supplementary Figure 1:**
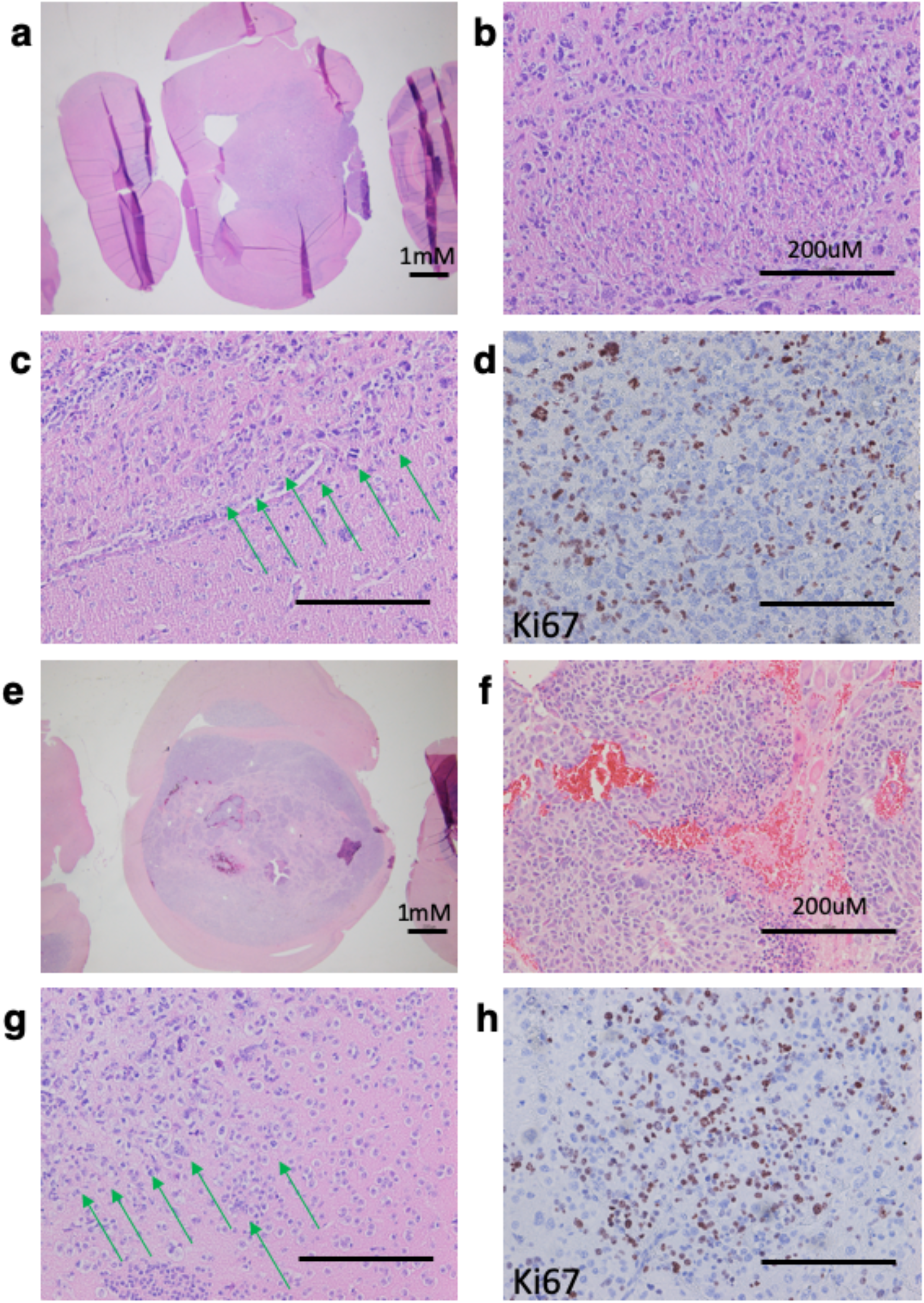
Histopathological staining of representative QPP gliomas arising spontaneously or implanted in mice, displaying the hallmark features of GBM. **a**. Representative hematoxylin and eosin (H & E)-stained whole-mount coronal section of brain with spontaneous QPP tumor at 1x magnification. **b**. Necrosis of the spontaneous tumor. **c**. Invasive infiltrating edge of the spontaneous tumor marked by green arrows. **d**. Ki67 staining within the spontaneous QPP glioma demonstrating the high number of active proliferating cells. **e**. Representative H & E-stained whole-mount coronal section of brain with implanted QPP tumor at 1x magnification (scale bar = 200 µM). **f**. Necrosis of the QPP implanted tumor. **g**. Invasive infiltrating edge of the implanted QPP tumor. **h**. Ki67 staining within the implanted QPP border to determine location of proliferating cells.

**Supplementary Figure 2:**
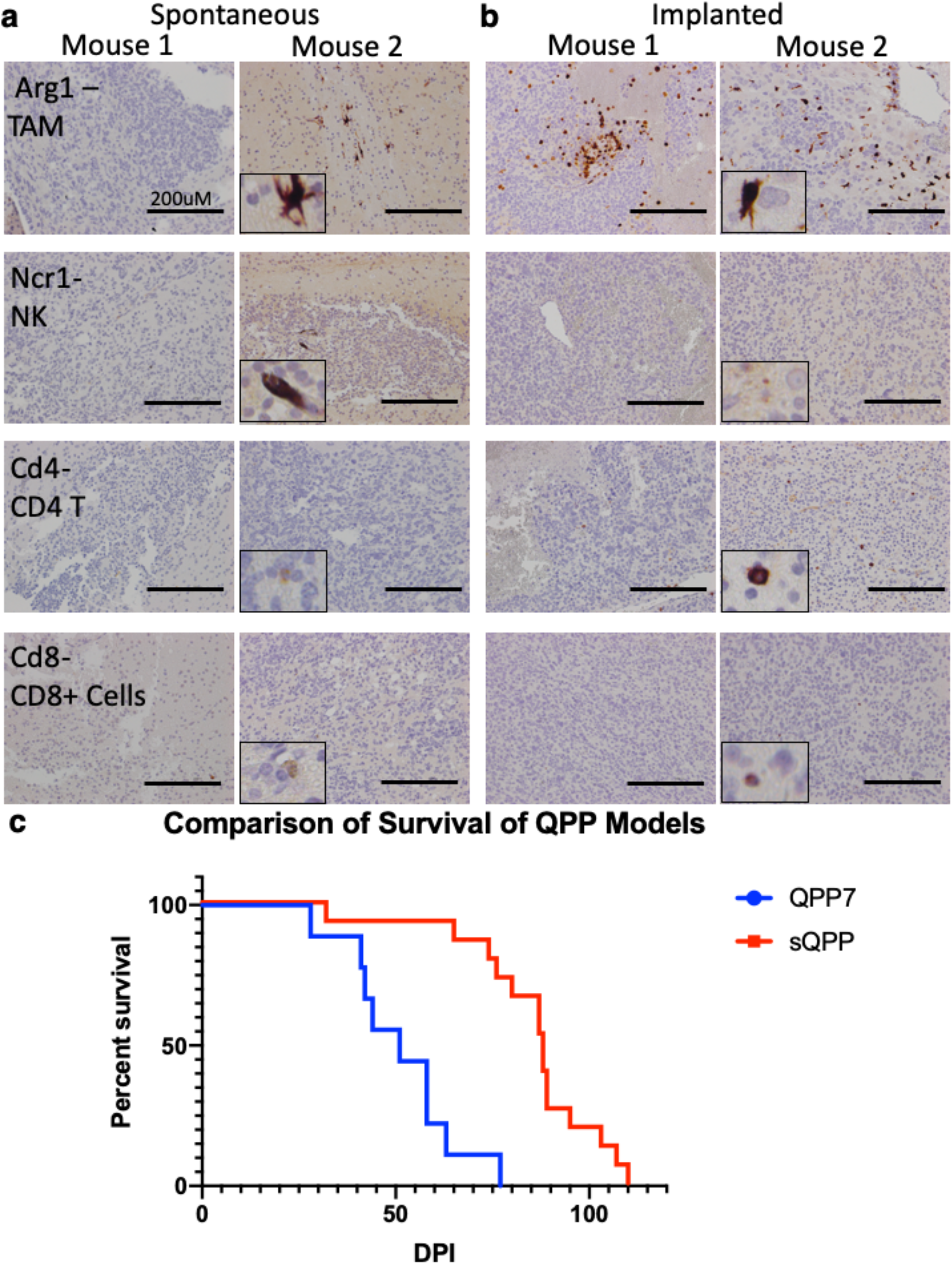
Kaplan-Meier Survival Curves for the QPP Model. **a**. Survival of n=15 QPP mixed background mice injected with 20uL subcutaneous tamoxifen on P7 and P8. **b**. Survival of n=9 C57Bl6/j mice implanted with 50,000 QPP7 cells into their striatum.

**Supplementary Figure 3:**
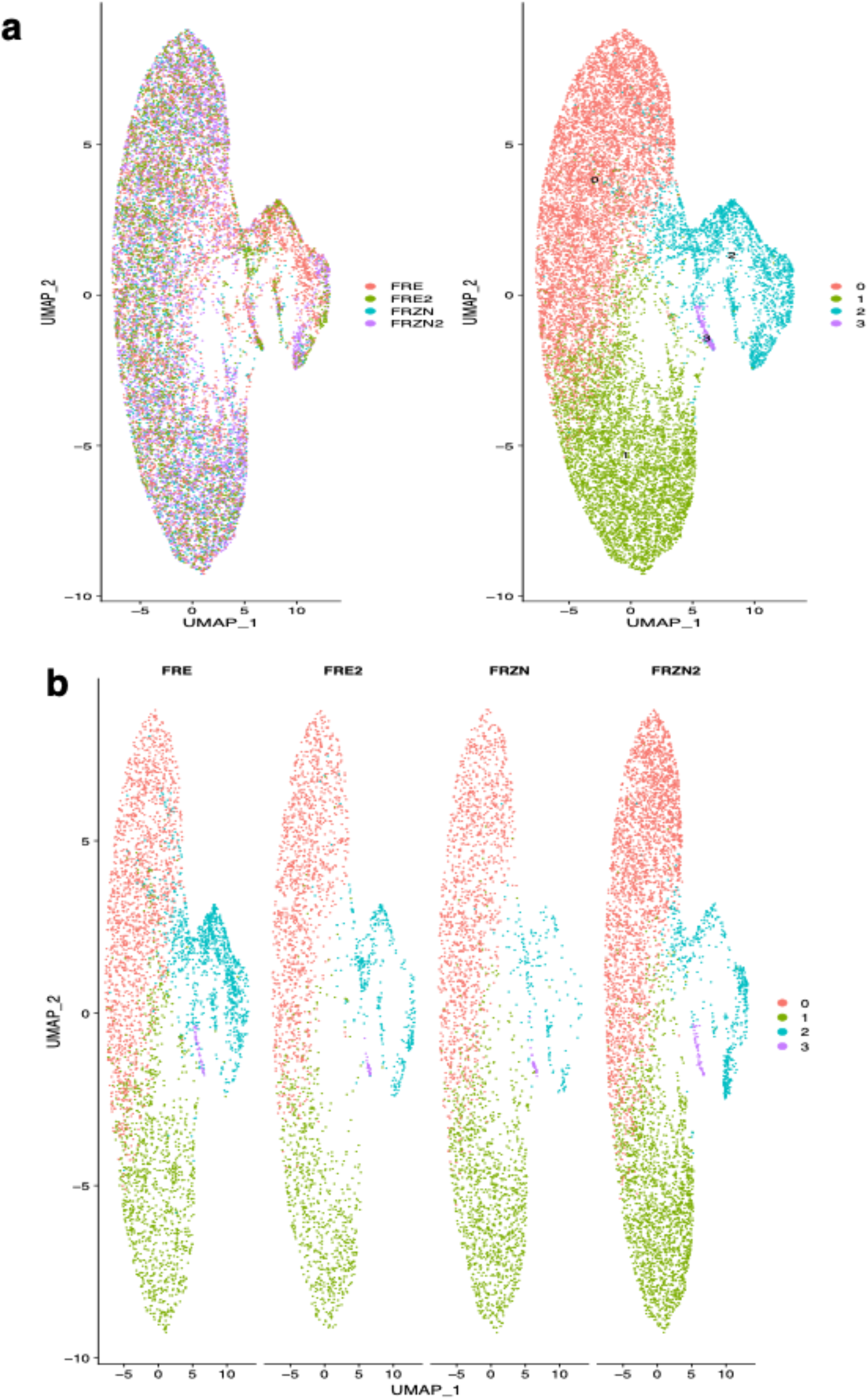
Quantification of Intratumoral 20x Fields for Immune Infiltrates. 3 separate fields from 5 independent mice were stained for either Cd3 (T-cells) or Cd11b (Myeloid cells) and quantified.

**Supplementary Figure 4:**
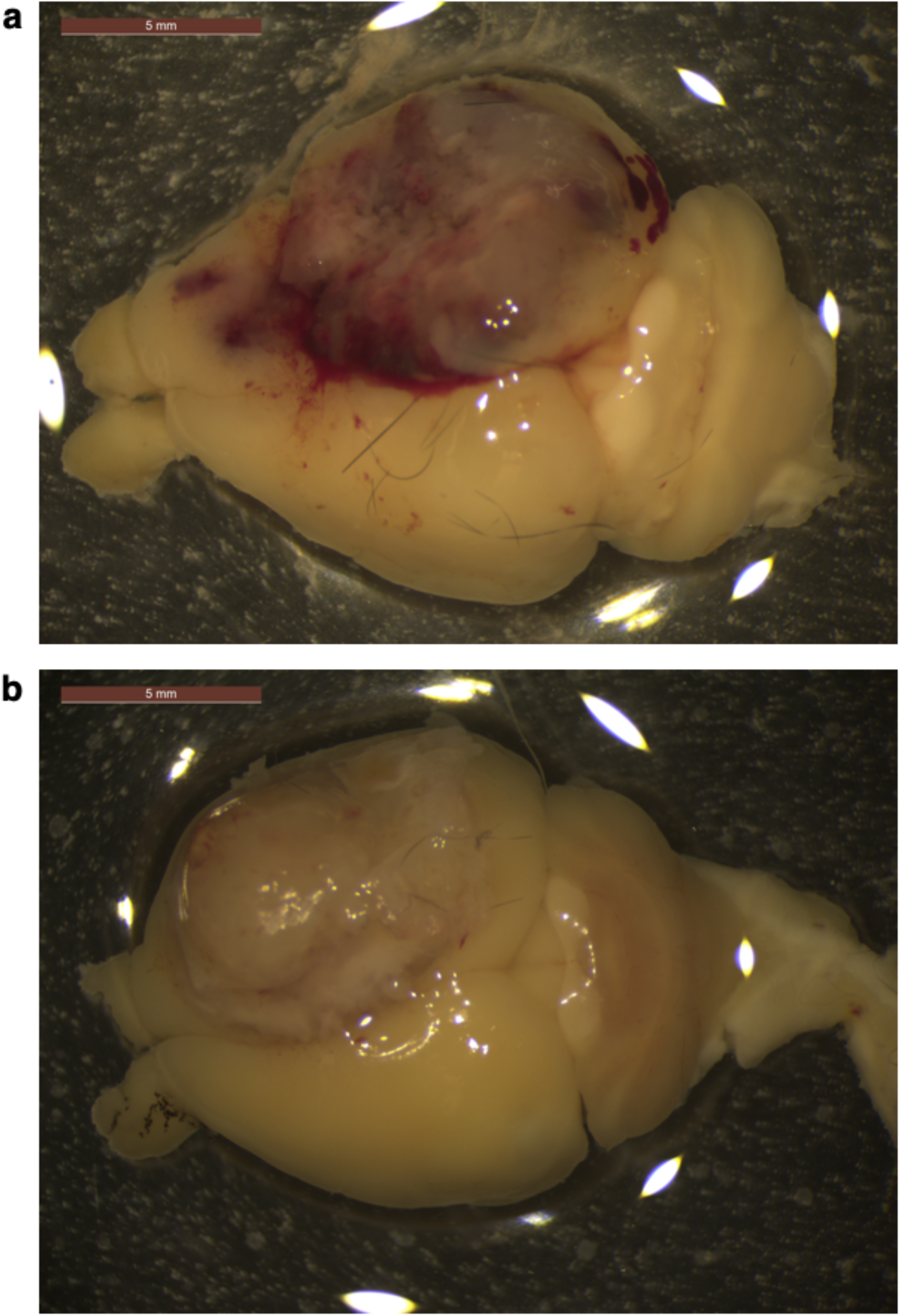
Comparison of Freshly Isolated Naïve Mouse Brains to Previously Frozen Naïve Mouse Brains. **a**. Aggregated clustering CD45+ cells of n=2 freshly isolated naïve mouse brains and n=2 previously frozen mouse brains. **b**. Demonstrates the contributions of individual mice to **Supplementary Figure 3a**.

**Supplementary Figure 5:**
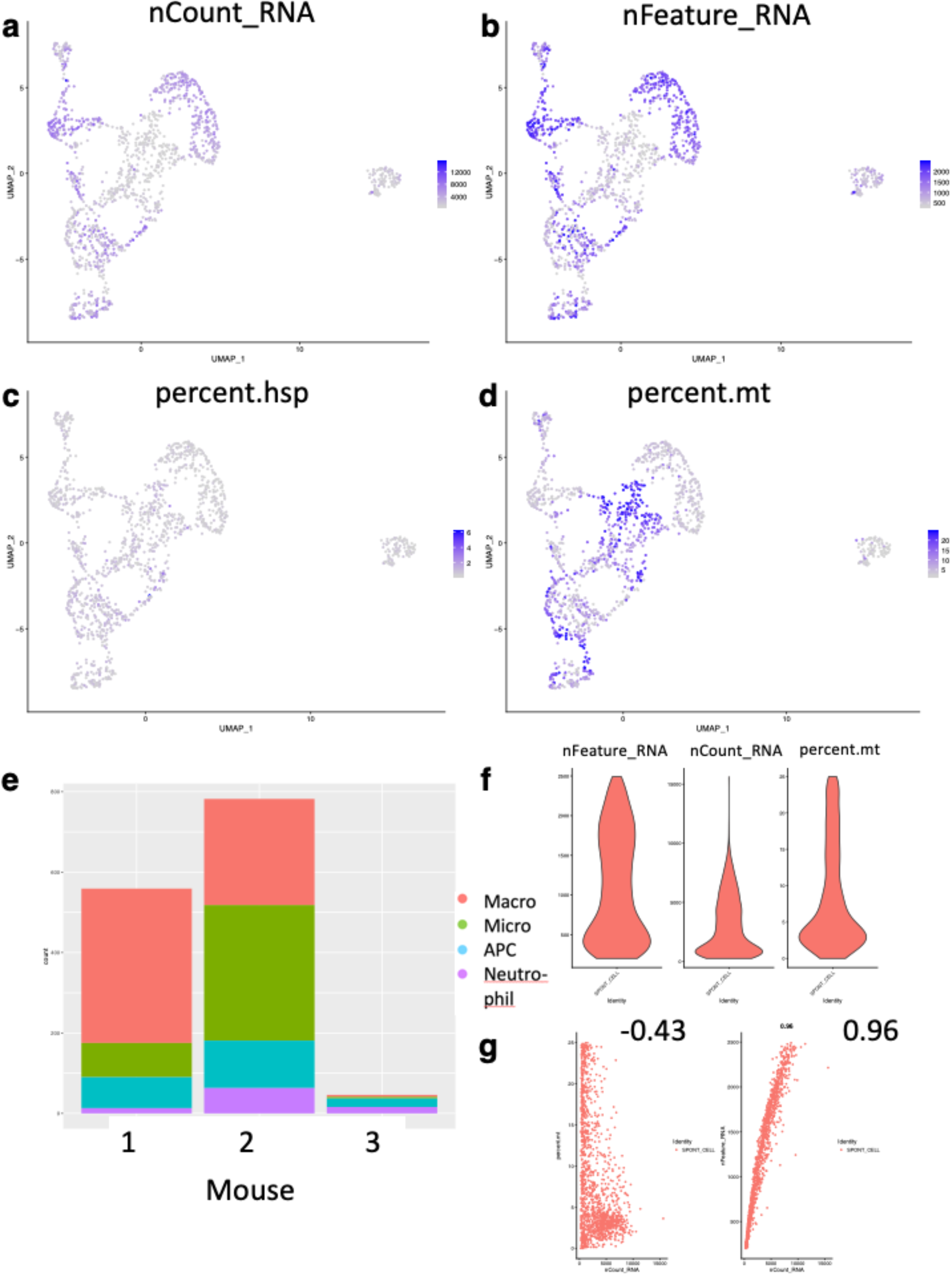
Immune Constituents of Spontaneous QPP. Demonstrative images of how clusters are labeled a given immune population. **a**. Shows aggregate of CD45+ immune infiltrates from n=3 spontaneous QPP tumors at moribund timepoint. **b**. Markers used to Identify the microglia and macrophage clusters and violin plots to show the specificity of given markers. **c**. Markers used to Identify the neutrophil cluster and violin plots to show the specificity of given markers. **d**. Markers used to Identify the APC cluster and violin plots to show the specificity of given markers. **e**. Markers to show that a population of T-cells exists with complementary violin plots. **f**. Markers to show that a population of NK-cells exists with complementary violin plots **g**. Markers to show that a population of B-cells exists with complementary violin plots

**Supplementary Figure 6:**
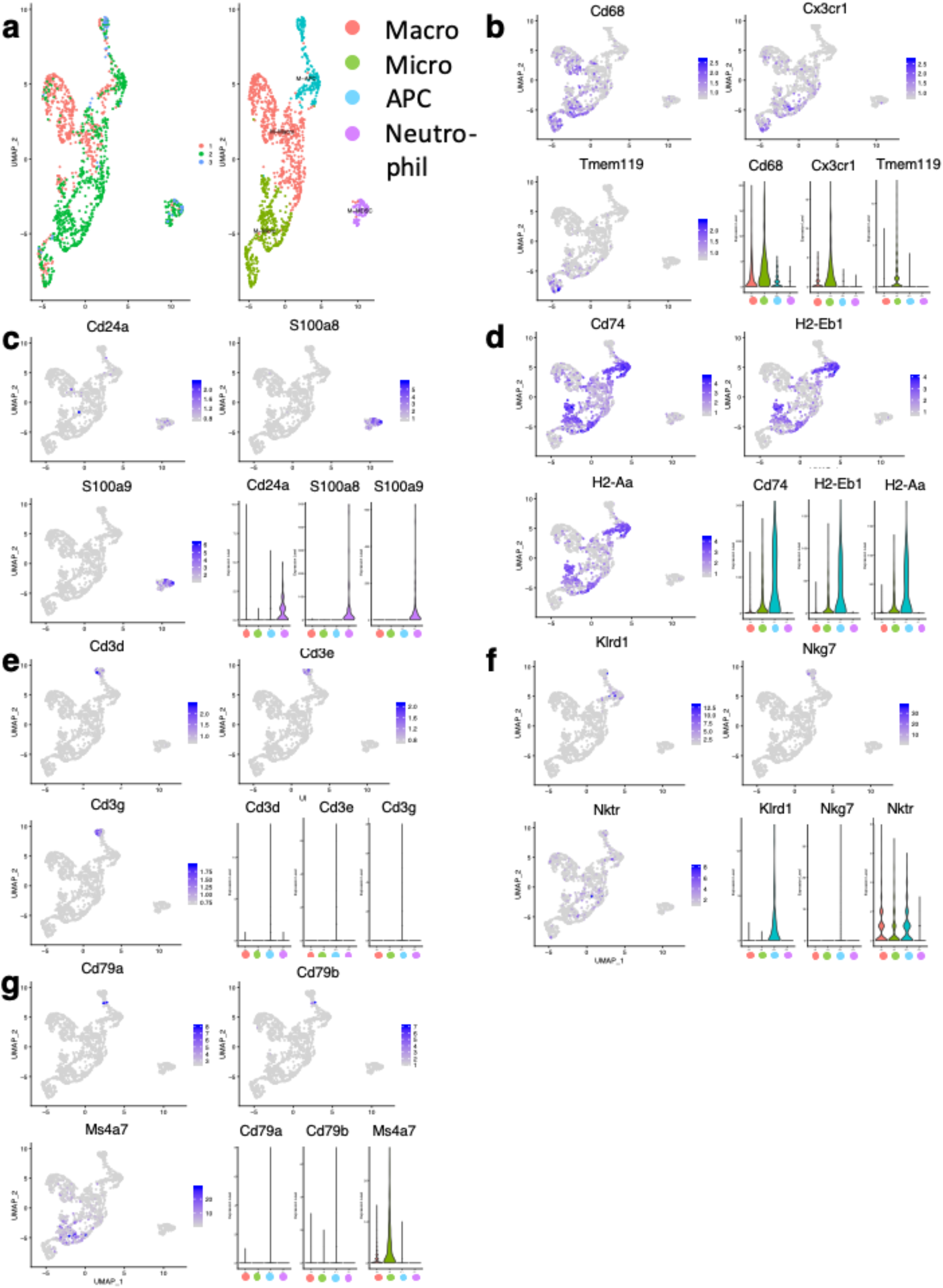
Immune Constituents of Implanted QPP. Demonstrative images of how clusters are labeled a given immune population. **a**. Shows aggregate of CD45+ immune infiltrates from n=3 Implanted QPP tumors at moribund timepoint. **b**. Markers used to Identify the microglia and macrophage cluster and violin plots to show the specificity of given markers. **c**. Markers used to Identify the neutrophil cluster and violin plots to show the specificity of given markers. **d**. Markers used to Identify the APC cluster and violin plots to show the specificity of given markers. **e**. Markers used to identify the lytic myeloid cluster and violin plots to show the specificity of given markers **f**. Markers to show that a population of NK-cells exists with complementary violin plots **g**. Markers to show that a population of B-cells exists with complementary violin plots **h**. Markers to show that a population of T-cells exists with complementary violin plots.

**Supplementary Figure 7:**
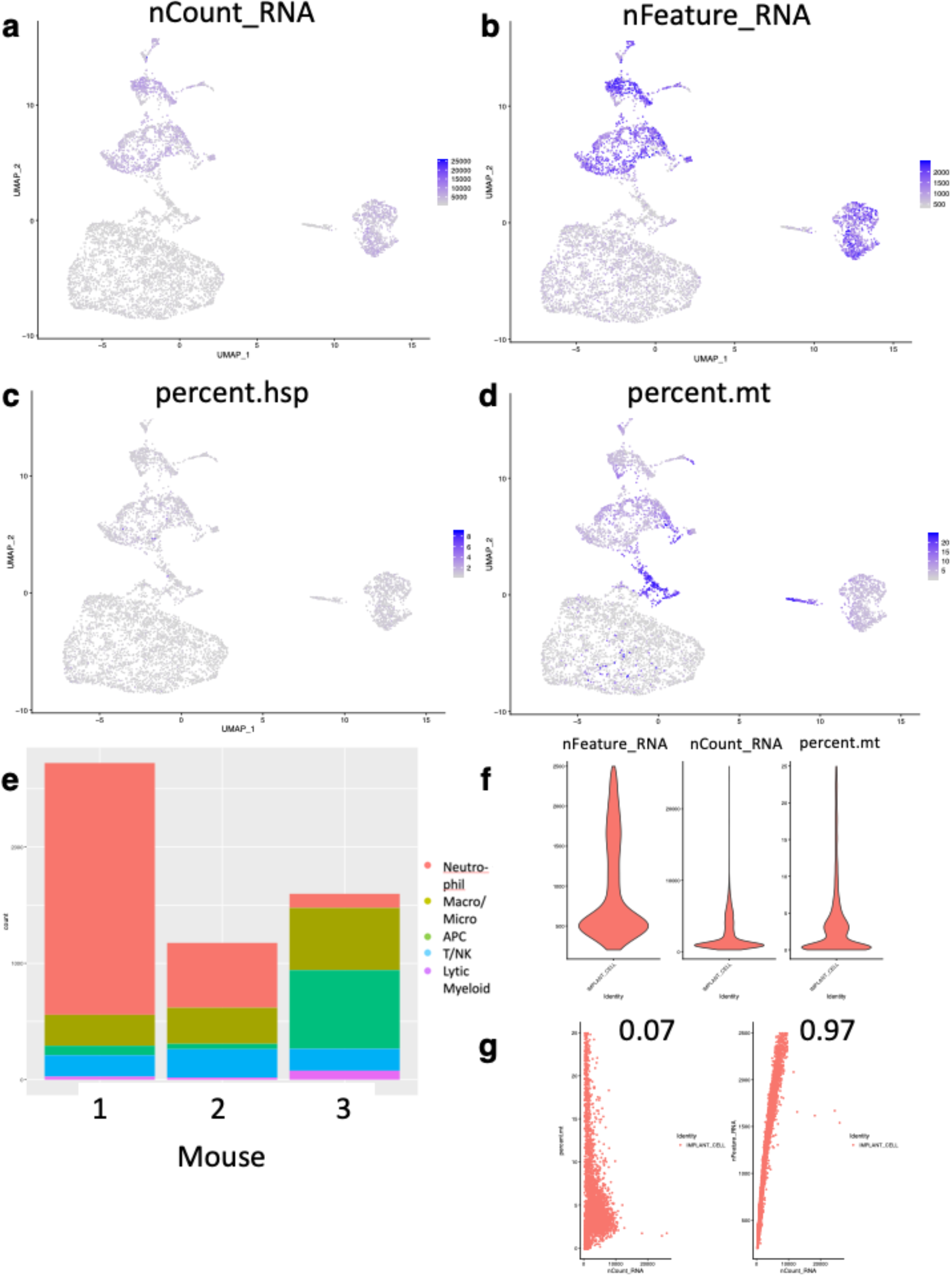
Comparison of QPP Immune Constituents. Demonstrative images of how clusters are labeled a given immune population. **a**. Shows aggregate of CD45+ immune infiltrates from n=3 spontaneous QPP tumors and n=3 Implanted QPP tumors at moribund timepoint. **b**. Markers used to identify neutrophil cluster and violin plots to show the specificity of given markers. **c**. Markers used to identify the microglia and macrophage cluster and violin plots to show the specificity of given markers. **d**. Markers used to identify the APC cluster and violin plots to show the specificity of given markers. **e**. Markers used to identify the lytic myeloid cluster and violin plots to show the specificity of given markers **f**. Markers to show that a population of NK-cells exists with complementary violin plots **g**. Markers to show that a population of B-cells exists with complementary violin plots **h**. Markers to show that a population of T-cells exists with complementary violin plots.

**Supplementary Figure 8:**
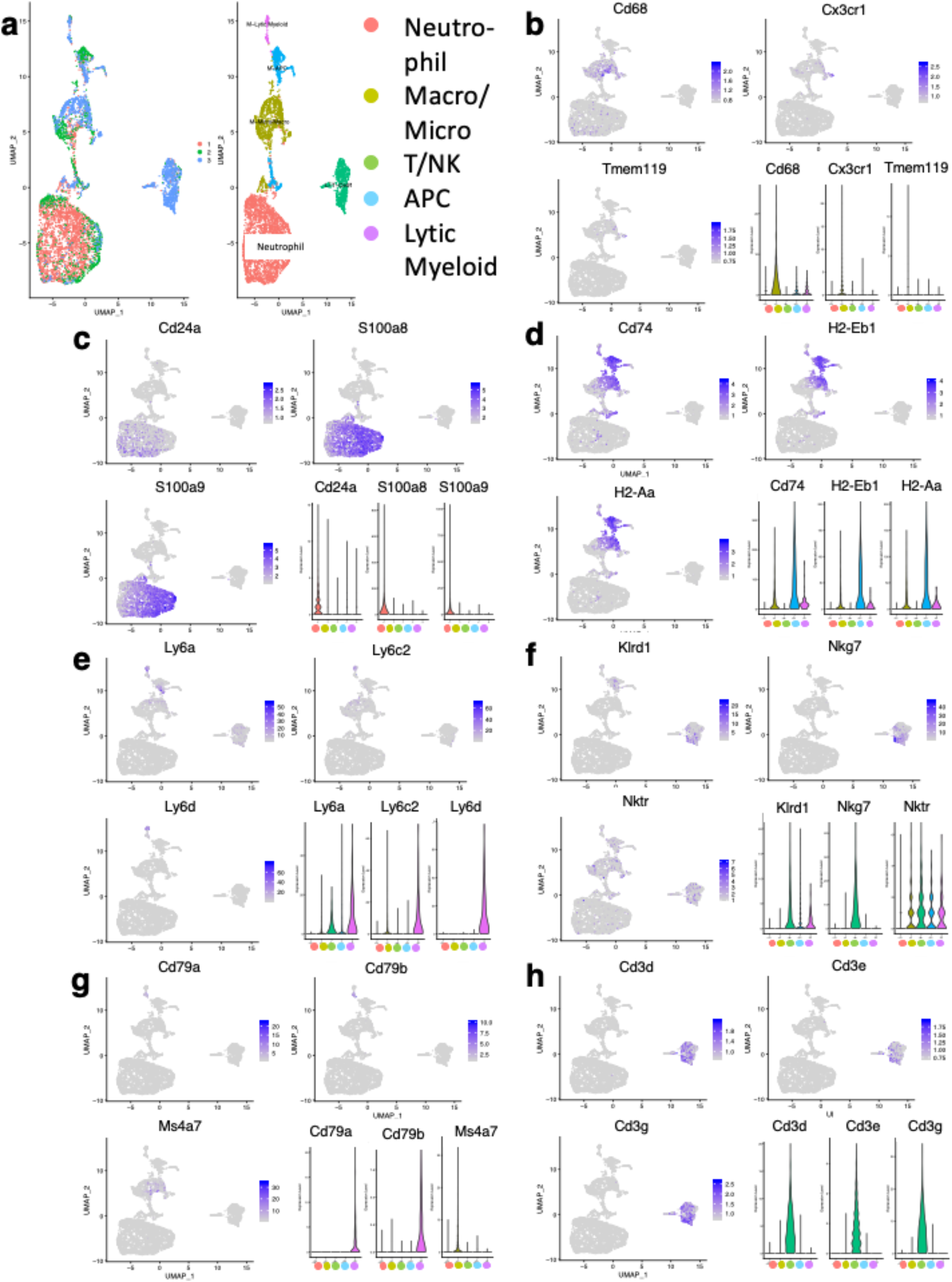
Heterogeneity between Individual Mice and Individual Patients. **a**. Demonstrates the contribution of the individual mice to the aggregate plot in **Figure 3a. b**. Demonstrates the contribution of the individual mice to the aggregate plot in **Figure 3b. c**. Demonstrates the contribution of the individual patients to the aggregate plot in **Figure 3d**.

**Supplementary Figure 9:**
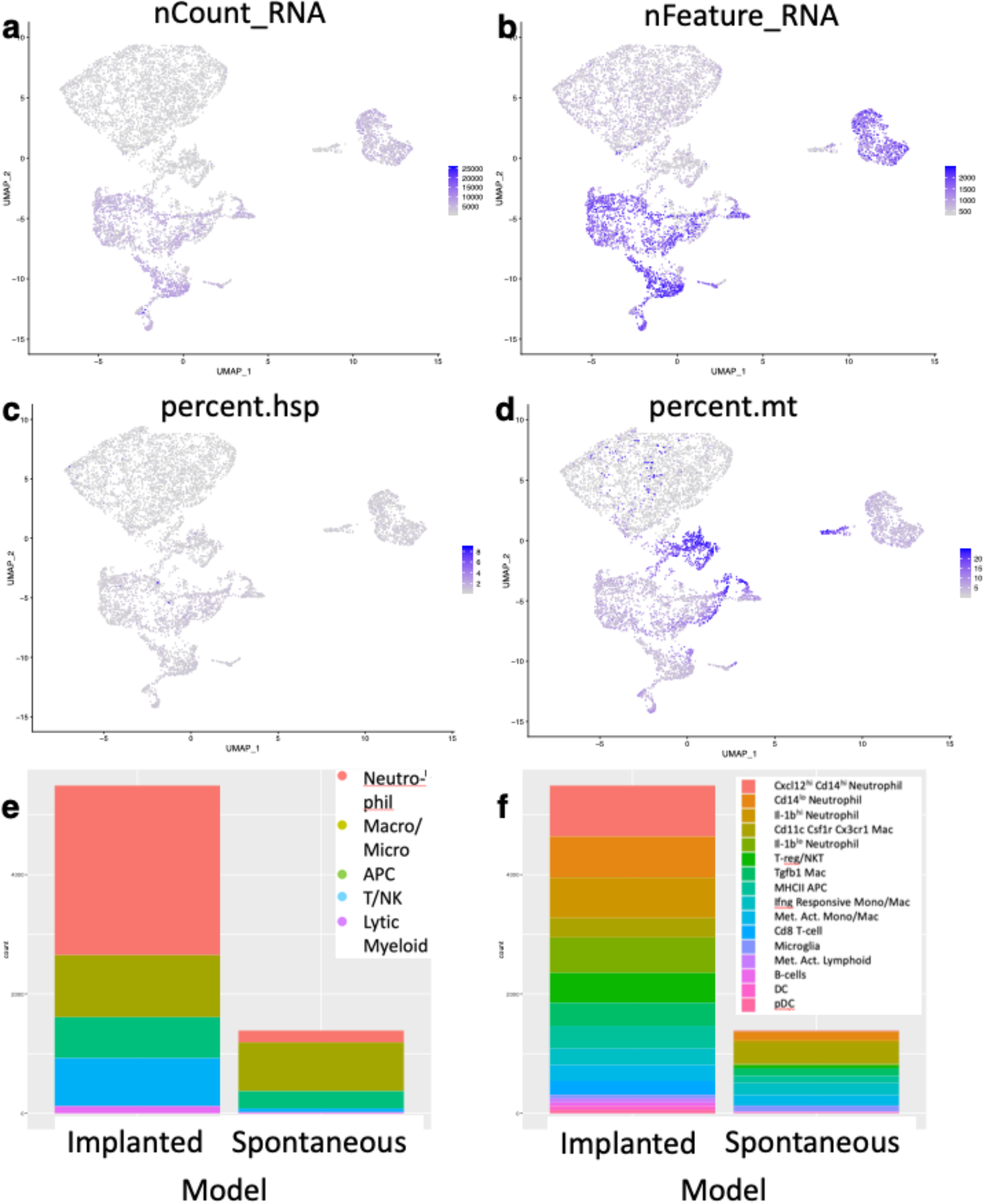
Immune Constituents of Human GBM. Demonstrative images of how clusters are labeled a given immune population. **a**. Shows aggregate of CD45+ immune infiltrates from n=10 glioma patients at surgical resection. **b**. Markers used to Identify the T-cell cluster and violin plots to show the specificity of given markers. **c**. Markers used to Identify the neutrophil cluster and violin plots to show the specificity of given markers. **d**. Markers used to Identify the NK cluster and violin plots to show the specificity of given markers. **e**. Markers used to identify the APC-like cluster and violin plots to show the specificity of given markers **f**. Markers to show that a population of microglia exists with complementary violin plots **g**. Markers to show that a population of macrophages exists with complementary violin plots **h**. Markers to show that a population of B-cells exists with complementary violin plots.

**Supplementary Figure 10:**
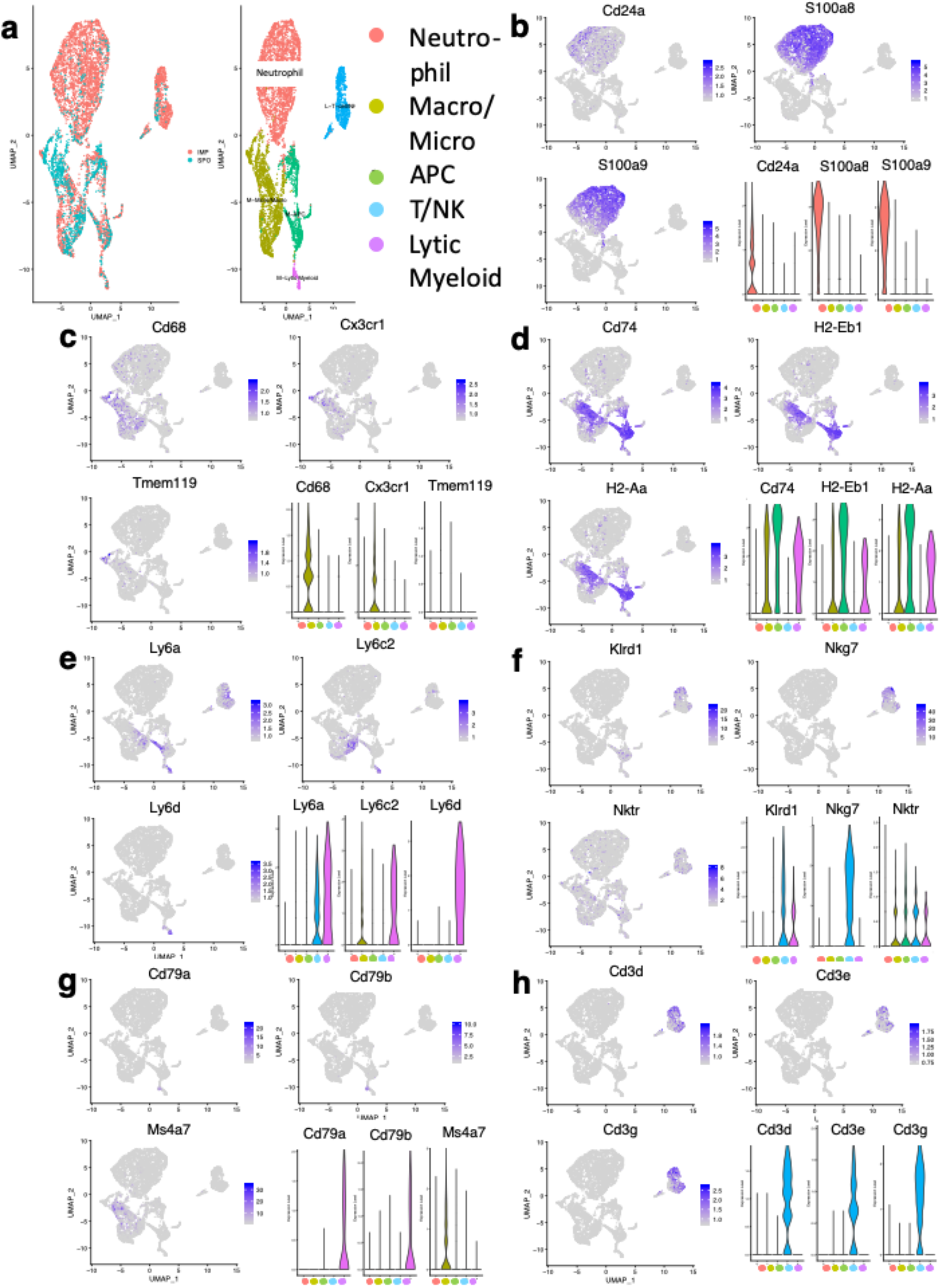
Immune Constituents of Human GBM Compared to Human LGG. **a**. Shows aggregate of CD45+ immune infiltrates from n=7 GBM patients at surgical resection compared to n=3 LGG patients at the same time point. **b**. Shows the GBM and LGG patients overlayed to better display the similarity and differences between the groups **c**. Markers used to Identify the upregulated myeloid cluster in LGG. **d**. Markers used to Identify the upregulated T cell cluster in LGG.

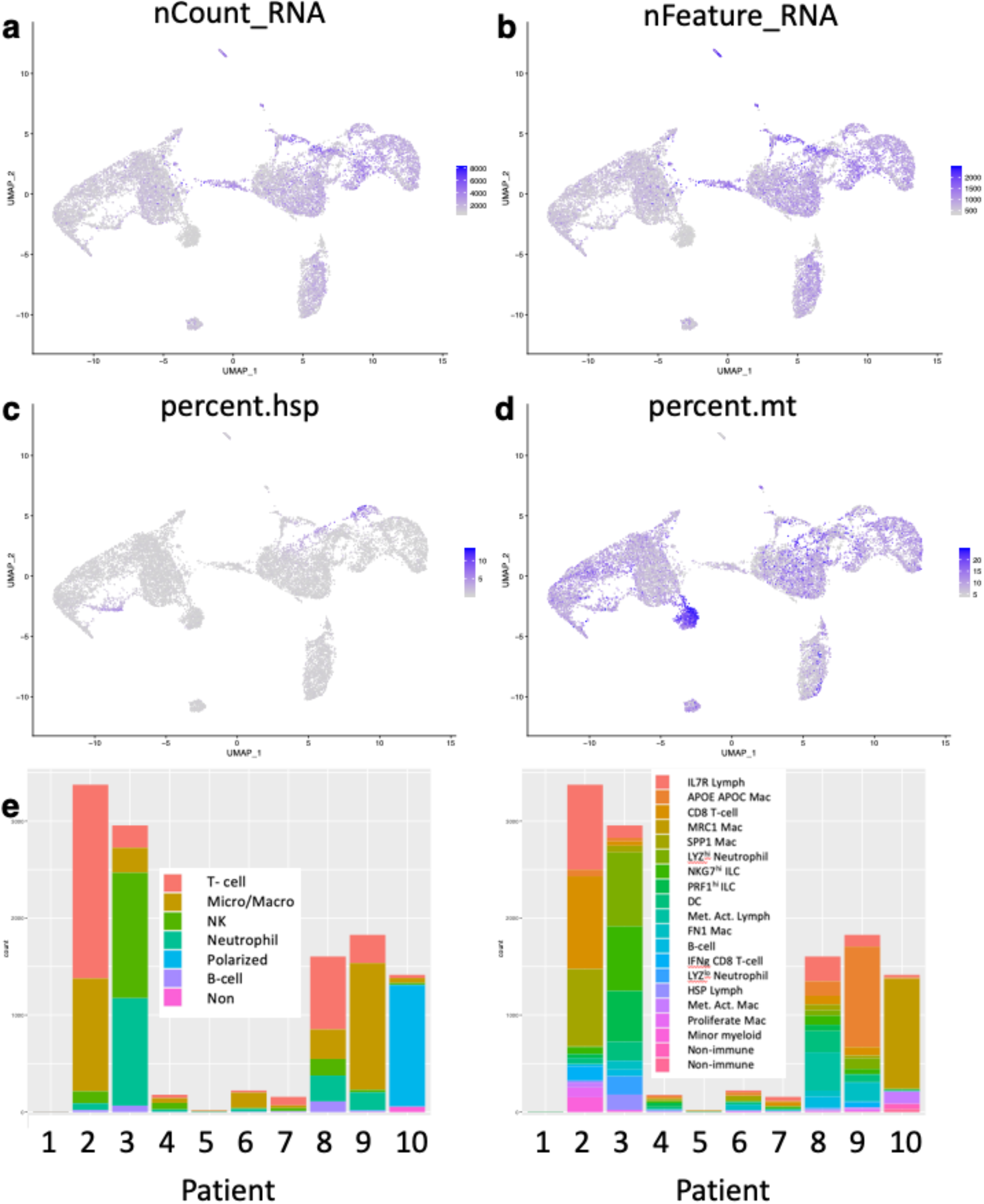

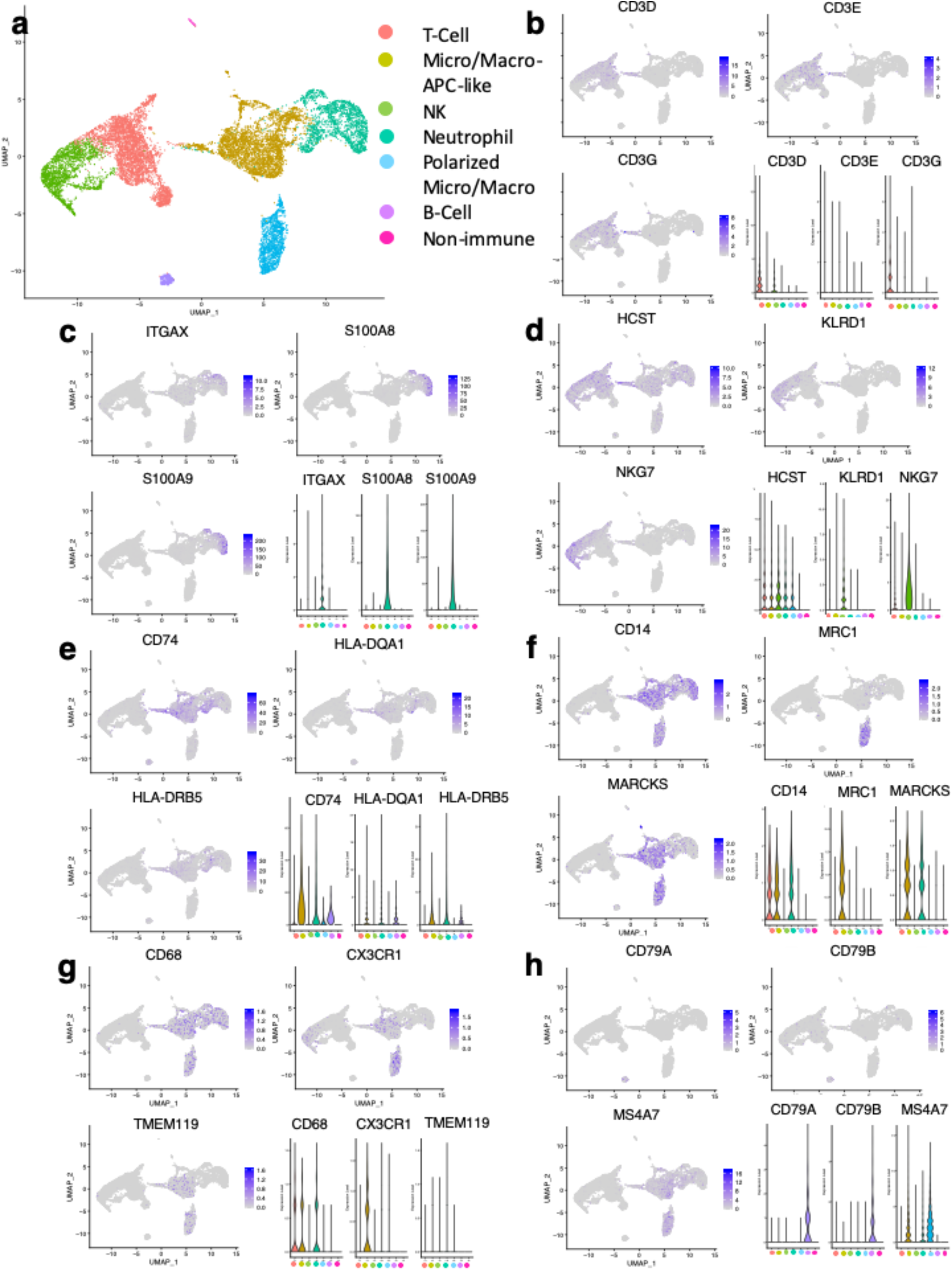

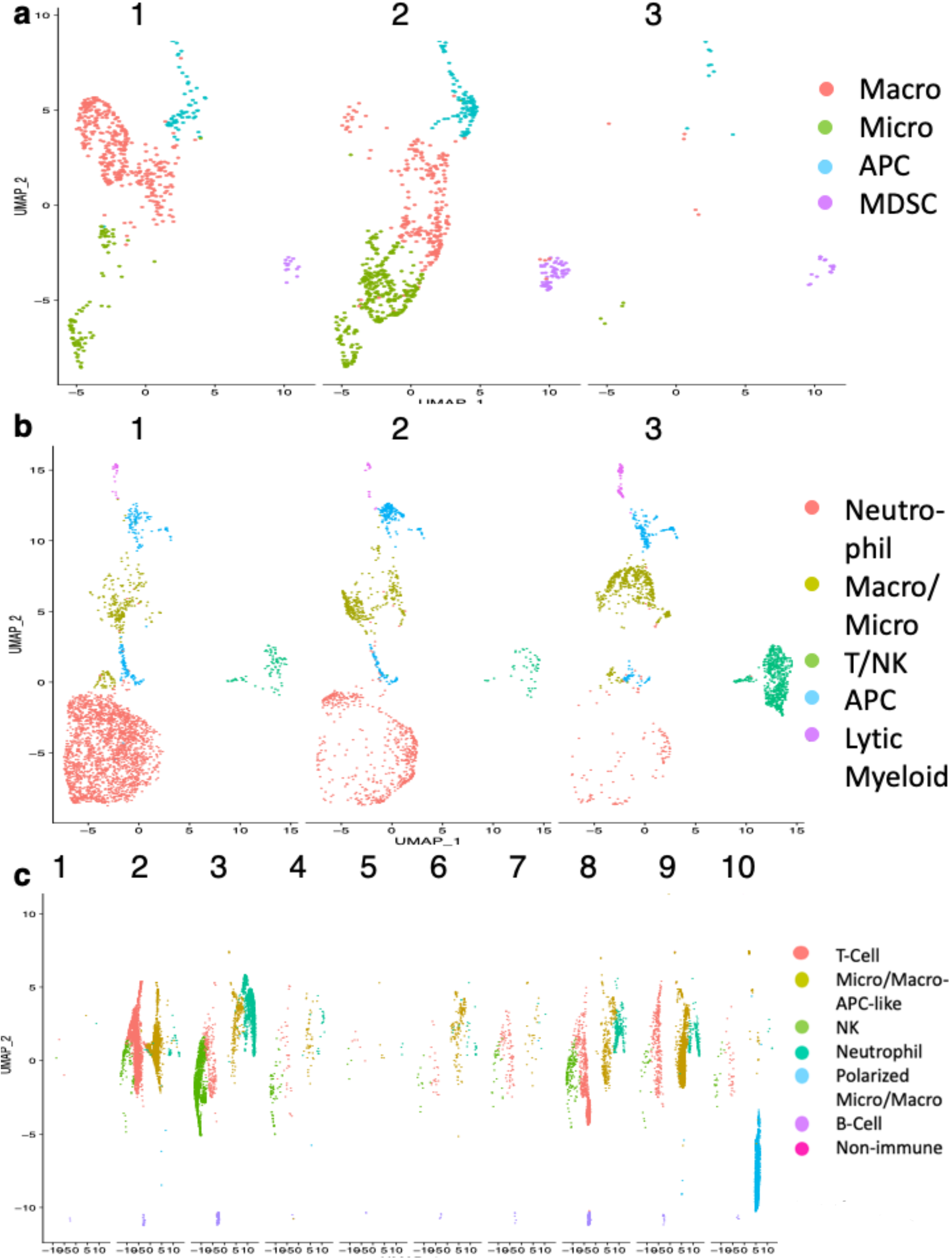

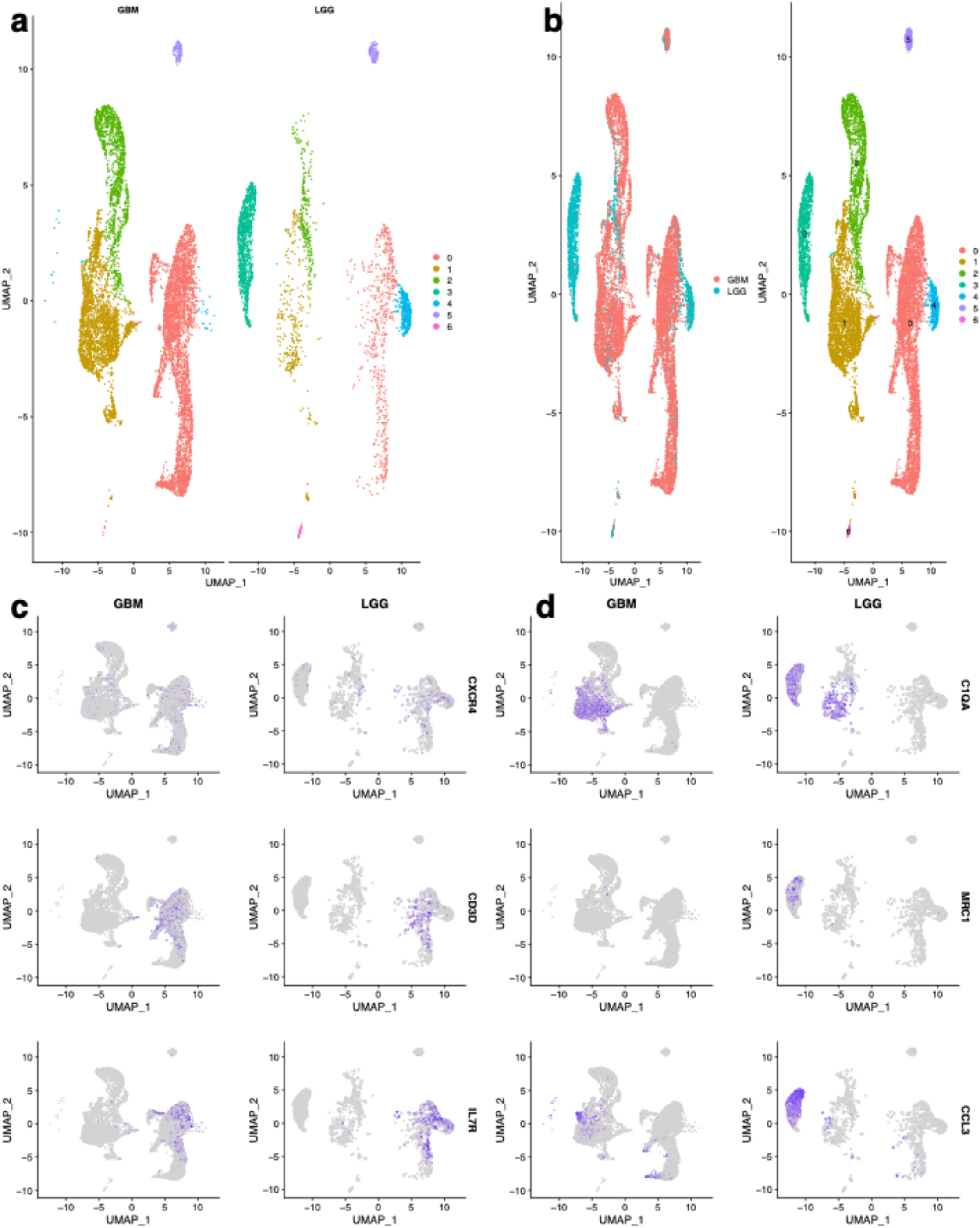

## REFERENCES

Bandopadhayay, P. et al. (2016) ‘MYB-QKI rearrangements in angiocentric glioma drive tumorigenicity through a tripartite mechanism’, Nature Genetics. Nature Publishing Group, 48(3), pp. 273–282. doi: 10.1038/ng.3500.

Brennan, C. W. et al. (2013) ‘The somatic genomic landscape of glioblastoma’, Cell, 155(2), p. 462. doi: 10.1016/j.cell.2013.09.034.

Butler, A. et al. (2018) ‘Integrating single-cell transcriptomic data across different conditions, technologies, and species’, Nature Biotechnology, 36(5), pp. 411–420. doi: 10.1038/nbt.4096.

Cerami, E., Gao, J., Dogrusoz, U., Gross, Benjamin E, et al. (2012) ‘The cBio cancer genomics portal: an open platform for exploring multidimensional cancer genomics data.’, Cancer discovery, 2(5), pp. 401–4. doi: 10.1158/2159-8290.CD-12-0095.

Cerami, E., Gao, J., Dogrusoz, U., Gross, Benjamin E., et al. (2012) ‘The cBio Cancer Genomics Portal: An open platform for exploring multidimensional cancer genomics data’, Cancer Discovery, 2(5), pp. 401–404. doi: 10.1158/2159-8290.CD-12-0095.

Chen, A. J. et al. (2012) ‘STAR RNA-binding protein Quaking suppresses cancer via stabilization of specific miRNA’, Genes and Development, 26(13), pp. 1459–1472. doi: 10.1101/gad.189001.112.

Chen, P. et al. (2019) ‘Symbiotic Macrophage-Glioma Cell Interactions Reveal Synthetic Lethality in PTEN-Null Glioma’, Cancer Cell. Elsevier Inc., 35(6), pp. 868-884.e6. doi: 10.1016/j.ccell.2019.05.003.

Chen, P. et al. (2020) ‘Circadian regulator CLOCK recruits immune suppressive microglia into the GBM tumor microenvironment’, 10(3), pp. 371–381. doi: 10.1158/2159-8290.CD-19-0400.Circadian.

Chen, S. (2020) ‘Gene Ontologies of scSeq Data’. GitHub. Available at: https://github.com/chansigit/scSnippet.

Chen, Z. and Hambardzumyan, D. (2018) ‘Immune Microenvironment in Glioblastoma Subtypes.’, Frontiers in immunology, 9(May), p. 1004. doi: 10.3389/fimmu.2018.01004.

Chongsathidkiet, P. et al. (2018) ‘Sequestration of T cells in bone marrow in the setting of glioblastoma and other intracranial tumors’, Nature Medicine. Springer US, 24(9), pp. 1459– 1468. doi: 10.1038/s41591-018-0135-2.

Friebel, E. et al. (2020) ‘Single-Cell Mapping of Human Brain Cancer Reveals Tumor-Specific Instruction of Tissue-Invading Leukocytes’, Cell, 181(7), pp. 1626-1642.e20. doi: 10.1016/j.cell.2020.04.055.

Gao, J. et al. (2013) ‘Integrative analysis of complex cancer genomics and clinical profiles using the {cBioPortal.}’, Sci Signal, 6(269), p. pl1. doi: 10.1126/scisignal.2004088.

Genoud, V. et al. (2018) ‘Responsiveness to anti-PD-1 and anti-CTLA-4 immune checkpoint blockade in SB28 and GL261 mouse glioma models’, OncoImmunology. Taylor & Francis, 7(12), pp. 1–10. doi: 10.1080/2162402X.2018.1501137.

De Groot, J. F. et al. (2018) ‘Window-of-opportunity clinical trial of a PD-1 inhibitor in patients with recurrent glioblastoma.’, Journal of Clinical Oncology. American Society of Clinical Oncology, 36(15_suppl), p. 2008. doi: 10.1200/JCO.2018.36.15_suppl.2008.

Hafemeister, C. and Satija, R. (2019) ‘Normalization and variance stabilization of single-cell RNA-seq data using regularized negative binomial regression’, bioRxiv, p. 576827. doi: 10.1101/576827.

Hussain, S. F. et al. (2006) ‘The role of human glioma-infiltrating microglia/macrophages in mediating antitumor immune responses1’, Neuro-Oncology, 8(3), pp. 261–279. doi: 10.1215/15228517-2006-008.

Johanns, T. M. et al. (2016) ‘Endogenous Neoantigen-specific CD8 T Cells Identified in Two Glioblastoma Models Using a Cancer Immunogenics Approach’, Cancer Immunology Research, 176(3), pp. 139–148. doi: 10.1016/j.physbeh.2017.03.040.

Kadić, E. et al. (2017) ‘Effect of cryopreservation on delineation of immune cell subpopulations in tumor specimens as determinated by multiparametric single cell mass cytometry analysis’, BMC Immunology, 18(1), pp. 1–15. doi: 10.1186/s12865-017-0192-1.

Klemm, F. et al. (2020) ‘Interrogation of the Microenvironmental Landscape in Brain Tumors Reveals Disease-Specific Alterations of Immune Cells’, Cell. Elsevier Inc., 181(7), pp. 1643-1660.e17. doi: 10.1016/j.cell.2020.05.007.

Mestas, J. and Hughes, C. C. W. (2004) ‘Of Mice and Not Men: Differences between Mouse and Human Immunology’, The Journal of Immunology, 172(5), pp. 2731–2738. doi: 10.4049/jimmunol.172.5.2731.

Miyai, M. et al. (2017) ‘Current trends in mouse models of glioblastoma’, Journal of Neuro-Oncology. Springer US, 135(3), pp. 423–432. doi: 10.1007/s11060-017-2626-2.

Ohlfest, J. R. et al. (2005) ‘Combinatorial antiangiogenic gene therapy by nonviral gene transfer using the Sleeping Beauty transposon causes tumor regression and improves survival in mice bearing intracranial human glioblastoma’, Molecular Therapy. The American Society of Gene Therapy, 12(5), pp. 778–788. doi: 10.1016/j.ymthe.2005.07.689.

Qaddoumi, I. et al. (2016) ‘Genetic alterations in uncommon low-grade neuroepithelial tumors: BRAF, FGFR1, and MYB mutations occur at high frequency and align with morphology’, Acta Neuropathologica. Springer Berlin Heidelberg, 131(6), pp. 833–845. doi: 10.1007/s00401-016-1539-z.

Shay, T. et al. (2013) ‘Conservation and divergence in the transcriptional programs of the human and mouse immune systems’, Proceedings of the National Academy of Sciences of the United States of America, 110(8), pp. 2946–2951. doi: 10.1073/pnas.1222738110.

Shingu, T. et al. (2017) ‘Qki deficiency maintains stemness of glioma stem cells in suboptimal environment by downregulating endolysosomal degradation’, Nature Genetics, 49(1), pp. 75– 86. doi: 10.1038/ng.3711.

Stuart, T. et al. (2019) ‘Comprehensive Integration of Single-Cell Data’, Cell. Elsevier Inc., 177(7), pp. 1888-1902.e21. doi: 10.1016/j.cell.2019.05.031.

Szatmári, T. et al. (2006) ‘Detailed characterization of the mouse glioma 261 tumor model for experimental glioblastoma therapy’, Cancer Science, 97(6), pp. 546–553. doi: 10.1111/j.1349-7006.2006.00208.x.

Verhaak, R. G. W. et al. (2010) ‘Integrated Genomic Analysis Identifies Clinically Relevant Subtypes of Glioblastoma Characterized by Abnormalities in PDGFRA, IDH1, EGFR, and NF1’, Cancer Cell, 17(1), pp. 98–110. doi: 10.1016/j.ccr.2009.12.020.

Woroniecka, K. et al. (2018) ‘T-cell exhaustion signatures vary with tumor type and are severe in glioblastoma’, Clinical Cancer Research, 24(17), pp. 4175–4186. doi: 10.1158/1078-0432.CCR-17-1846.

Yue, F. et al. (2014) ‘A comparative encyclopedia of DNA elements in the mouse genome’, Nature. Nature Publishing Group, 515(7527), pp. 355–364. doi: 10.1038/nature13992.

